# Sequence-based genome-wide association studies reveal the polygenic architecture of *Varroa destructor* resistance in Western honey bees *Apis mellifera*

**DOI:** 10.1101/2024.02.16.580755

**Authors:** Sonia E. Eynard, Fanny Mondet, Benjamin Basso, Olivier Bouchez, Yves Le Conte, Benjamin Dainat, Axel Decourtye, Lucie Genestout, Matthieu Guichard, François Guillaume, Emmanuelle Labarthe, Barbara Locke, Rachid Mahla, Joachim de Miranda, Markus Neuditschko, Florence Phocas, Kamila Tabet, Alain Vignal, Bertrand Servin

**Author notes:** These authors contributed equally to the work (Sonia E. Eynard, Fanny Mondet). These authors jointly supervised the work (Alain Vignal, Bertrand Servin).

## Abstract

Honey bees, *Apis mellifera*, have experienced the full impacts of globalisation, including the recent invasion by the parasitic mite *Varroa destructor* which has become one of the main causes of colony losses worldwide. Despite its lethal effects, some colonies have developed defence strategies conferring colony resistance and, assuming non-null heritability, selective breeding of naturally resistant bees could be a sustainable way to fight infestations. Here we report on the largest genome-wide association study performed on honey bees to understand the genetic basis of multiple phenotypes linked to varroa resistance. This study was performed on whole genome sequencing of more than 1,500 colonies belonging to different ancestries and combined in a meta-analysis. Results show that varroa resistance is polygenic. A total of 60 genetic markers were identified as having a significant impact in at least one of the tested populations pinpointing several regions of the honey bee genome. Our results also support strategies for genomic selection in honey bee breeding.

## Introduction

The honey bee, *Apis mellifera*, is a crucial contributor to sustainable food production [1]. However, for the past two decades beekeepers have been experiencing dramatic colony losses [2, 3]. Such losses are not sustainable for the beekeepers and the agroecosystems relying on services provided by honey bees. Extensive research has shown that honey bees are threatened by multiple factors: both abiotic factors with the loss of natural resources and the impact of pesticides due to agriculture intensification, and biotic factors with the infection by a diversity of pests and parasites that impair their survival [4]. Among biotic factors, the ectoparasite *Varroa destructor* is currently considered as the main threat to honey bee health and beekeeping worldwide [5]. In most regions of the world, colony losses have dramatically increased since its introduction in *Apis mellifera* populations in the early 80s [5, 6]. Originating from Asia, where a stable host-parasite relationship exists with its historical host *Apis cerana*, varroa now infests most *Apis mellifera* colonies worldwide. Varroa infests multiple compartments of the honey bee colony: it reproduces in the brood, feeds on adult honey bee haemolymph and fat body and favours virus infections [5, 7, 8]. Combined, these effects on individual bees lead to colony collapse within a few months if no actions are applied to control mite infestations [9]. To date, managing varroa infestation presents many constraints that offer beekeepers only a few unsustainable solutions to fight the deadly mite [7, 10].

However, since the beginning of the 1990’s, colonies naturally surviving varroa infestation without treatment have been observed in several regions of the world and raised hope for beekeepers to overcome the problems caused by varroa infestation. In these surviving colonies, undergoing beekeeping activities, honey bees often display behavioural and physiological defences against the varroa parasite. These collective responses are expected to contribute to the limitation of parasite population growth and provide colonies with social immunity [11], a sustainable long-term adaptation to counter the immense damage caused by varroa. However, thus far such long term adaptation is not always reached, even in colonies expressing resistance traits (e.g. colonies in Gotland, Sweden [12, 13]). The defence repertoire against varroa includes hygiene behaviour targeted towards varroa parasitised brood cells, in the form of varroa sensitive hygiene (VSH [14]) or recapping behaviour [15, 16, 17], and mechanisms suppressing mite reproduction (SMR [18]). Expression of these different traits lead to an increase in mite non reproduction within brood cells (MNR [15, 18], also called Decreased Mite Reproduction DMR [19]. This triggers lower mite population growth and infestation [10]. Such mechanisms have gained a major interest within the beekeeping sector and interest is growing to help decipher the genetic mechanisms underlying varroa resistance in the honey bee.

As for common livestock species, the most obvious way to induce varroa resistance into the honey bee population appears to be through selection and spreading of the most resistant lines. Since the late 90’s efforts have been put into the selective breeding of such resistant honey bee lines [20, 21]. However, selecting for complex traits in honey bees has been hindered by features that strongly differentiate honey bees from other typical livestock species. Due to the social nature of honey bees, many phenotypic traits of interest to beekeeping (including varroa resistance traits) are expressed at the group (i.e. colony) level and thus can not be addressed by classical GWAS approaches where phenotypes are determined at the individual level (e.g. individual bees, in this case). In addition, many of these traits, are linked to group behaviour and are thus difficult to phenotype and can display rather low repeatability and heritability [22, 23, 24, 25, 26]. At the breeding level, managing honey bee reproduction is difficult due to polyandry [27] (queen bees mate with multiple males) and to the fact that sex determination is governed by an haplodiploid mechanism at a single locus [28, 29] at which diploid homozygosity is effectively lethal [30], limiting drastically the possibility of inbreeding. Finally, honey bee populations bred for beekeeping encompass an important genetic diversity, with strong regional clustering, which impedes the identification of genetic markers that are valid outside the population where they are identified. Developing molecular tools to assess the genetic make-up of honey bee colonies could help in honey bee breeding by coping with some of these issues. To list a few possibilities, genomic-enabled prediction of resistance traits could reduce the amount of complex phenotyping required by identifying promising colonies early in life and genetic assessment of colonies could help identifying the genetic group of colonies [31] and allow to design crosses limiting inbreeding in selection programs.

In addition to their use in selection programs, molecular tools can help to identify pathways involved in a specific mechanism by performing genome-wide association studies, assessing the statistical effects of polymorphisms on the variability of complex phenotypes. This offers opportunities to develop new approaches that take into account the specific genetic determinism of honey bees and to open avenues for genomic selection on traits such as varroa resistance. However, tools to perform genetic association studies are so far not tailored to encompass such genetic characteristics, which limits the power of genomic studies performed on honey bees and their transferability into breeding tools. Some markers of interest have been identified (see [16] for a review) but most genomic studies performed on honey bee traits so far were built on a limited number of samples (10 to 200 hundred individuals or colonies [32, 33, 34, 35, 36, 37, 38, 39], restricting the power of the analyses to come out as a general breeding tool. To date, the use of such markers has been limited, mostly due to a lack of easily accessible genotyping tools, leaving beekeepers with very limited access to varroa resistant stock.

In this study, we took advantage of the unique situation represented by France. Geographical crossroad, the French territory has the advantage to present a large variety of landscapes, environments and ecosystems, where honey bee populations with different genetic background coexist, together with a large variety of hybrid colonies [40]. To overcome the limitations in sample size, we performed one of the largest genomic study applied to honey bees, with the phenotyping and complete genome sequencing of more than 1,500 colonies. Using uniquely tailored genetic and genomic tools, such as queen genotype reconstruction from pool sequence data [31] and GWAS and meta-GWAS analysis [41, 42], we investigated the genetic bases of three major traits linked to varroa resistance: overall varroa infestation of the colony, mite non reproduction (MNR) and recapping of varroa infested brood cells. We identified multiple genetic markers of interest, spread out on the whole genome and heterogeneous across the different populations. This large-scale effort provides a new understanding of the genetic mechanism underlying honey bee resistance to its main parasite, the *Varroa destructor* mite.

## Results

### Genetic and phenotypic diversity of honey bee colonies

Using allele frequencies estimated from pool sequence data for the 1513 sampled honey bee colonies, we identified three groups of different genetic ancestries: 703 colonies were identified as having more than 80% *Apis mellifera ligustica & carnica* genetic background, 407 having more than 80% *Apis mellifera mellifera* genetic background and 382 as hybrids (Fig. 1). An additional 21 colonies were found to be of pure *Apis mellifera caucasia* ancestry, but due to this reduced sample size of this category they were not analysed further.

**Fig. 1:**
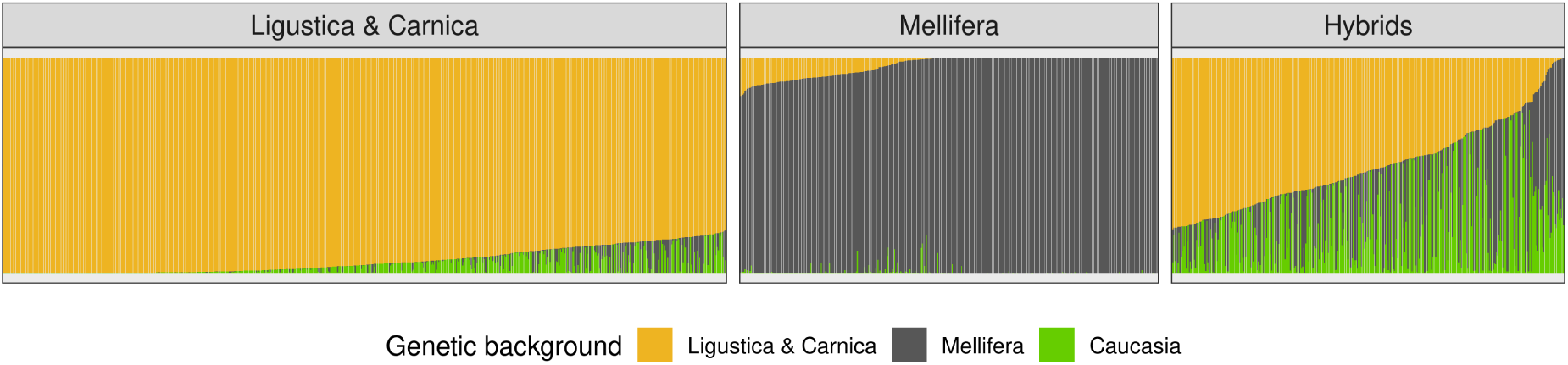
Genetic background for each colony, per group. Proportion of the three main genetic background for each of the group analysed in our study *Apis mellifera ligustica & carnica*, *Apis mellifera mellifera* and hybrids.

For the purpose of this study we collected six phenotypes on these colonies. Out of the six phenotypes initially available four were associated with varroa infestation (on the adult bees: phoretic infestation rate *v_pho* and varroa mitochondrial sequence reads *v_mito*, inside the brood: brood infestation rate *v_brood* and overall in the colony: varroa load *v_load*). They were highly positively correlated with each other and drove the first dimension of the principal component analysis (PCA) across all colonies (Fig. 2), and also within each group (supplementary figure S1), with about 60% of the variance explained in each case. Therefore, coordinates of colonies on the first component of the PCA were used as the varroa infestation phenotype in the genome wide association study (GWAS) (*varroa_inf*). The two remaining phenotypes were expected to be linked to resistance to varroa infestation, either through mechanisms repressing varroa reproduction, and thus varroa population growth, within the colony (so called mite non reproduction) or through the cleaning of brood cells infested by varroa. Mite non reproduction (*MNR*) was slightly positively correlated with the recapping of infested brood cells, as expected under the assumption that *recap* contributes to overall *MNR*. Both *MNR* and *recap* were slightly negatively correlated with varroa infestation. This observation is consistent with mechanisms being linked to a reduction in varroa infestation within the colony. *MNR* and *recap* contributed both to the second dimension of the PCA, explaining about 20% of the variance. They are separated on the third axis of the PCA, which explains about 12% of the variance (Fig. 2). GWAS were perfomed on each of these three phenotypes (*varroa_inf*, *MNR* and *recap*).

**Fig. 2:**
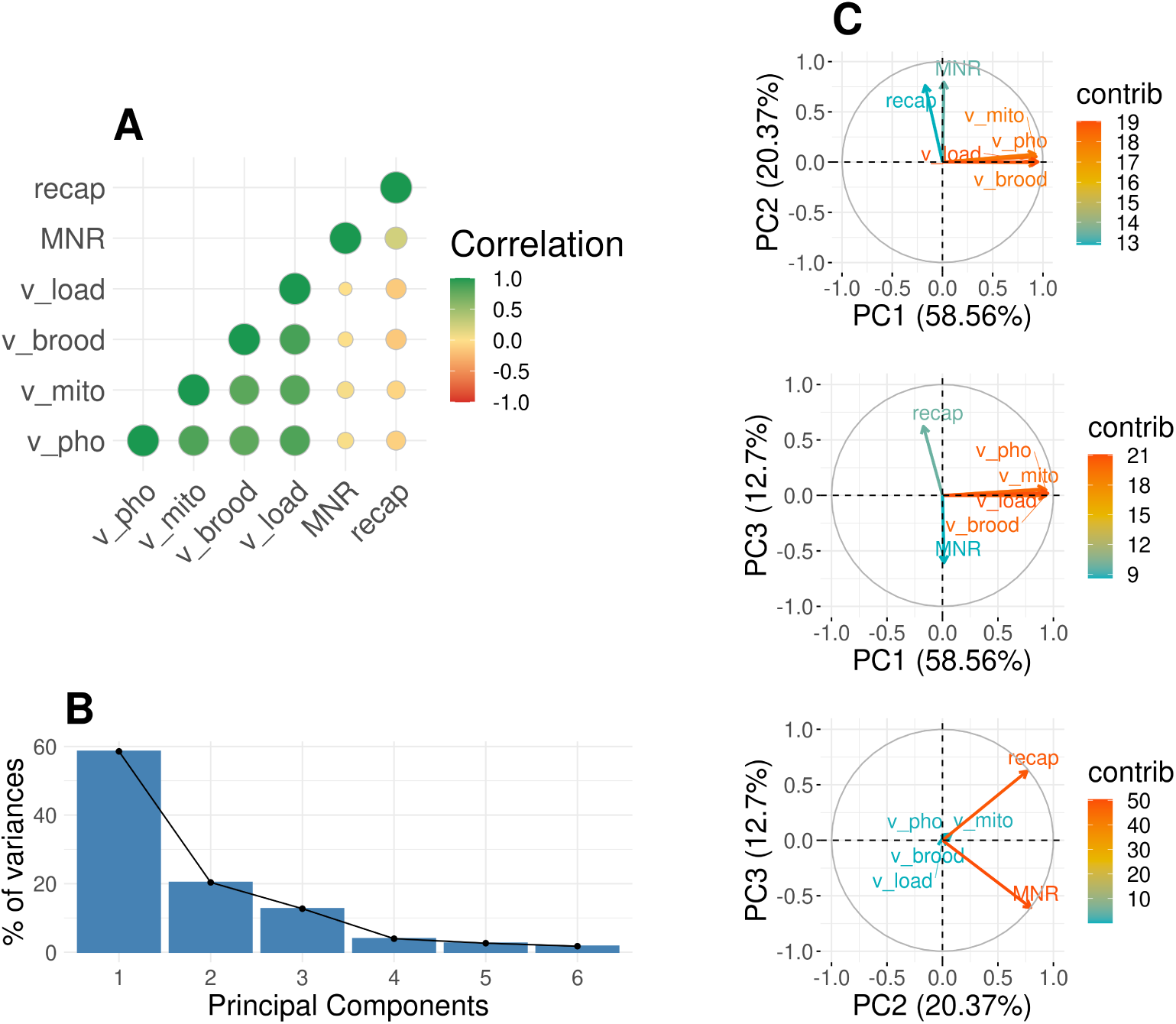
Correlation and principal component analysis. Description of the correlation between phenotypes and principal component analysis. **(A)** gives the correlation between our original phenotypes. **(B)** summarises the percentage of variance explained by each of the principal component analysis from axis 1 to 5. **(C)** shows our phenotype on principal component analysis for axis 1, 2 and 3, the colour gives the contribution of each variable to the axis, the closer to red, the higher. The correlation and PCA estimates are based on an analysis of the whole dataset, including all colonies from the three groups.

### Meta-analysis of varroa resistance

We aim to identify markers significantly associated with our traits of interest across the three main genetic backgrounds found in Europe. Within population we used standard linear mixed model equations, implemented in GEMMA [43], significant markers were identified relative to their local false discovery and false sign rates estimates from adaptive shrinkage [44]. Population level analyses were combined into a meta-analysis, using two Bayesian methods: a co-ancestry specific program MANTRA [41] and the more generalist Mash [42], to increase power to detect significant markers across the three genetic backgrounds.

#### Associated variants

The individual GWAS for each genetic background group and each phenotype allowed to identify 8 regions (n SNPs=9).

In details:

i. For *varroa_inf*, we found one variant (1:10080627:C>T) in *A. m. ligustica & carnica*, with a positive alternative allele effect, and one (4:11665460:G>A) in the hybrid group, with a negative alternative allele effect (a detailed list of the single nucleotide polymorphisms, SNPs, can be found in supplementary table SR1).
ii. For *recap* we identified four variants significant in *A. m. mellifera*, two of which the alternative alleles had negative effects on the trait (2:2729874:T>C and 3:1059430:T>C) and two having positive effects (4:4327611:G>A, 15:8485332:G>A). In *A. m. ligustica & carnica* a region, with two SNPS, both of which alternative alleles had positive effects (2:12025610:A>G, 2:12025647:A>G) were found. Finally in the hybrid group, one variant for which the alternative allele had a negative effect on the trait (13:9483955:C>T) was found.

Out of the 9 SNPs found significant in one trait for one group, eight fell inside genes of the honey bee annotation [45] and one fell 7kb upstream of its closest gene (for more details see supplementary table SR1).

The meta-analysis permitted to identify 51 regions, containing 56 significant SNPs across the three traits of interest. From these 14 regions (n SNPs=14) were significant for *varroa_inf*, 14 regions (n SNPs=15) for *MNR* and 23 regions (n SNPs=27) for *recap*. They distributed across the whole genome, on almost every chromosome (a detailed list of the SNPs can be found in supplementary table SR2).

In details:

i. For the *varroa_inf* trait: the 14 SNPs having a significant effect on the trait were distributed on the chromosomes 7 (n=4), 1, 5 and 8 (n=2) and 4, 6, 11 and 12 (n=1). Thirteen of these SNPs were considered significant in the MANTRA meta-analysis with log10(BF) ranging from 7.59 to 5.09 and one was significant in the mash analysis with log10(BF)=1.16. For six of these variants, the alternative allele had a positive effect on the trait, in at least one of the group (5:9190579:A>G, 7:5772089:A>T, 7:6738985:T>A, 7:11806658:G>A, 8:9799408:C>T and 12:10734707:A>G), for seven alternative allele had a negative effect on the trait, in at least one of the group (1:20960056:C>T, 1:25184394:C>T, 4:11665460:G>A, 5:75369:C>T, 6:10450971:C>T, 7:5762037:T>C and 8:2468335:C>T) and for one the effect depended on the group (11:9369229:T>C). Eight SNPs fell inside genes found in the honey bee annotation, six of them are intronic variation, one leads to a change in 5’ UTR and one in 3’ UTR. Three fell downstream, within 54kb of their closest genes. The remaining three fell up upstream, within 11kb of their closest genes.
ii. For *MNR*, the 15 SNPs having a significant effect on the trait were distributed on the chromosomes 1 (n=4), 8, 10 and 12 (n=2) and 2, 3, 5, 11 and 15 (n=1). All these SNPs were considered significant in the MANTRA meta-analysis with log10(BF) ranging from 5.12 to 6.44. In three of these variants, the alternative allele had a positive effect on the trait, in at least one of the group (2:4437645:G>A, 12:10153855:A>G and 15:4853529:C>T), for seven it had a negative effect on the trait, in at least one of the group (1:16327085:C>T, 1:21374478:G>A, 1:24201224:C>T, 3:6206342:C>T, 8:1150346:C>T, 11:9527267:G>A and 12:136634:G>C) and for five the effect depended on the group (1:2891204:G>A, 5:2008472:A>C, 8:9557205:C>T, 10:5359169:T>A and 10:5359173:C>T). Out of the 15 significant SNPs, 12 fell inside genes, 10 are intronic variation, one into 3’ UTR and one causes a missense variation. The three remaining SNPs fell within 17kb downstream of their closest genes.
iii. Finally, for the *recap* trait SNPs were distributed on the chromosomes 2 and 7 (n=4), 1, 5 and 14 (n=3), 4 and 15 (n=2) and 3, 8, 9, 10, 11 and 16 (n=1). From these SNPs 26 were considered significant based on their log10(BF) values for the MANTRA meta-analysis with log10(BF) ranging from 5.01 to 7.61 and 6 SNPs add log10(BF) values from mash higher than one, their log10(BF) ranged between 1.41 and 2.53. Differently from the other trait five SNPs were found significant in both MANTRA and mash meta-analysis for *recap*. Their log10(BF_MANTRA) ranged between 5.03 and 7.61 and their log10(BF_mash) between 1.41 and 2.53. They were located on the chromosomes 2 (n=2), 3, 11 and 15. For eight of these variants, the alternative allele had a positive effect on the trait, in at least one of the group (1:7448807:A>T, 1:7448811:T>C, 4:7321246:T>A, 4:7321247:G>T, 14:6686131:A>G, 14:8481541:A>G, 15:2081876:A>G and 15:8485332:G>A), for 18 the alternative allele had a negative effect on the trait, in at least one of the group (1:15280956:G>A, 2:2729874:T>C, 2:8350714:G>A, 2:12025610:A>G, 2:16060868:G>A, 3:1059430:T>C, 5:6736534:T>C, 5:6761414:T>A, 5:8737386:G>A, 7:7028040:G>A, 7:7051965:A>G, 7:7078376:C>T, 8:1551638:C>T, 9:11564671:A>C, 10:2026877:C>G, 11:14369154:G>C, 14:3782741:G>A and 16:1812909:C>T) and for one the effect depended on the group (7:8466948:A>G). For this trait, 18 variants fell inside genes with 16 intronic variation, one in a region coding for long non coding RNA and one in 5’ UTR, being also a missense variant. Three SNPs fell between 52 and 102kb downstream of their closest genes. Six SNPs fell between 5 and 129kb upstream of their closest genes (details are available in supplementary tables SR2 and SR3).

We identified chromosome regions smaller than 1Mb, sharing significant SNPs between multiple traits. Between *varroa_inf* and *MNR* we identified two regions on chromosome 1 (20.9-21.4Mb and 24.2-25.2Mb), a region on chromosome 8 (9.5-9.8Mb), a region on chromosome 11 (9.3-9.6Mb) and a region on chromosome 12 (10.1-10.8Mb) having significant SNPs for both traits. Between *varroa_inf* and *recap* we identified a region on chromosome 5 (8.7-9.2Mb), a region on chromosome 7 (6.7-7.1Mb) and a region on chromosome 8 (1.5-2.5Mb). Finally between *MNR* and *recap* we identified a region on chromosome 8 (1.1-1.6Mb) (Fig. 3, for the details of each region see supplementary table SR4).

**Fig. 3:**
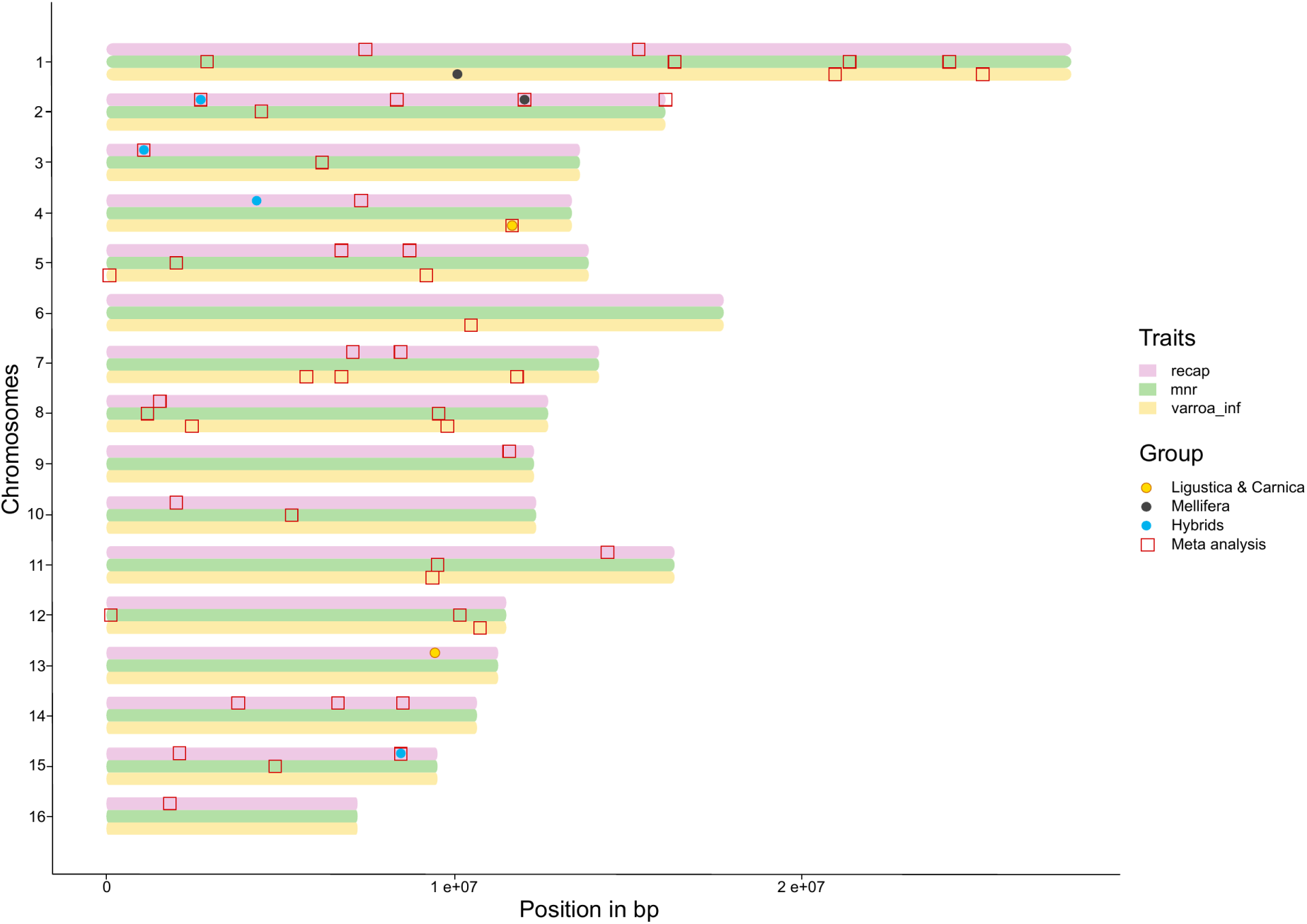
Overlap between traits across significant regions. Position of each of the significant SNPs on their chromosomes. A coloured dot means that this position has been identified in either *A. m. ligustica & carnica*, *A. m. mellifera* or hybrids. A red square means that it has been identified in the meta analysis. The colour bars represent the phenotypes of interest *varroa_inf*, *MNR* and *recap*. This figure allows to see overlapping window containing significant marker across the phenotypes.

#### Heterogeneity of effects

We analysed three genetic groups of colonies in this study, with two groups having relatively pure genetic backgrounds, corresponding to the two main lines of honey bee subspecies in Europe, with the third group consisting of varying degrees of hybridization between these two main groups. The Fst, measure of population differentiation, between groups were 0.26, between *A. m. mellifera* and *A. m. ligustica & carnica*, 0.20 between *A. m. mellifera* and hybrids and 0.08 between *A. m. ligustica & carnica* and hybrids.

As detailed earlier, from the meta analysis we observed 17 variants (n *varroa_inf* =6, n *MNR*=3 and n *recap*=8) for which alternative allele had significant positive effects for at least one of groups, 32 (n *varroa_inf* =7, n *MNR*=7 and n *recap*=18) having significant negative effects for at least one of the group, and seven (n *varroa_inf* =1, n *MNR*=5 and n *recap*=1) having divergent effect depending on the group (supplementary table SR3). Correlations between the effects of significant SNPs between different groups were null therefore showing great heterogeneity in SNPs having an effect, their impact on the trait of interest and their magnitude across the different genetic groups, especially for the *MNR* trait.

About 80% of the SNPs identified significant fell into intronic regions, there was no differences between the annotation of the significant SNPs on the genome and the annotation of all tested SNPs. Five SNPs were found significant in both individual GWAS and meta-analysis, one for *varroa_inf* identified significant for the hybrid group, and four for *recap*, three for *A. m. mellifera* and one for *A. m. ligustica & carnica*. They were located on chromosome 2 (n=2), 3 (n=1), 4 (n=1) and 15 (n=1). They all located inside the genes, LOC102655235, LOC410853 (chromosome 2), LOC409402 (chromosome 3), LOC408787 (chromosome 4) and LOC726948 (chromosome 15).

#### Example of associations

As an example of a region associated with two traits, we focus on the region between 9.5 and 9.8Mb on the chromosome 8 where two SNPs appear significant in the meta analysis, one for *MNR* at 9,557,205 bp and one for *varroa_inf* at 9,799,408 bp. In this region we identified a couple of SNPs in close vicinity to the gene Ecr, ecdysone receptor (Fig. 4). In addition, we also identified a SNP, located at 9,696,277bp so within Ecr, in high linkage disequilibrium (LD) with the significant SNP 8:2468335:C>T also found in the meta-analysis for *varroa_inf* (supplementary table SR6).

**Fig. 4:**
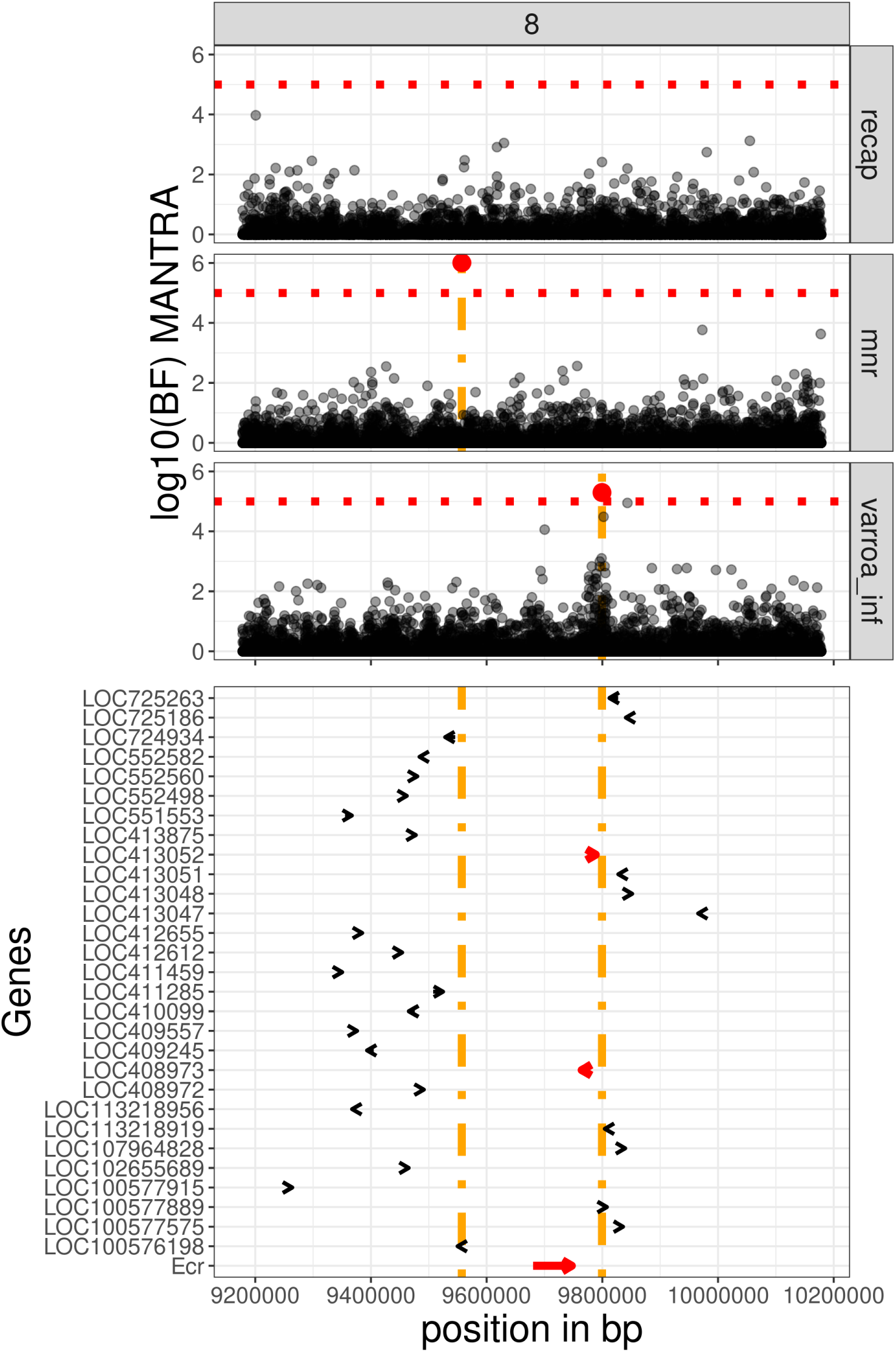
Region on chromosome 8. From top to bottom we represented results from genome wide association studies for chromosome 8, region going from 9.2Mb to 10.2Mb, for *varroa_inf*, *MNR* and *recap*. We represented log10 bayes factor for MANTRA, with the significant threshold as a dotted red line, under which we listed the genes identified within this region of chromosome 8. The orange lines represent the positions of the significant markers, the genes falling in the region between these markers are highlighted in red.

Next, we turn to chromosomes 15, where a significant SNP for *recap* was found at position 2,081,876. Nearby we also observed two suggestive SNPs, close to the significant threshold set for this analysis, in positions 2,021,142 and 2,081,914 bp. The markers 15:2081876:A>G and 15:2081914:A>G were in full linkage disequilibrium in *A. m. ligustica & carnica* and in the hybrid group and in high linkage disequilibrium (*r*^2^ > 0.8) in *A. m. mellifera* and they fell within the same haplotype block of 1.6kbp (as identified by Wragg et al. (2022) [40], supplementary table SR7). This haplotype block did not seem to contain annotated genes. The marker 15:2021142:C>T fell in a short haplotype block (0.175kbp) containing the gene LOC413200 which has been identified as putative immune related gene by Ryabov et al. (2014) [46]. Interestingly they were located within less than 1Mbp downstream from a group of eight genes coding for odorant binding proteins. For the first SNP (15:2021142:C>T), alternative allele had a negative effect for all groups whereas the second (15:2081876:A>G), mostly significant in *A. m. ligustica & carnica*, the alternative allele had a positive effect for all groups (Fig. 5). Marker effects estimated with GEMMA lack precision, whereas the ash and mash methods apply a strong shrinkage to the estimates. Going from individual analysis to meta-GWAS improved the power to detect associations and improve our ability to estimate markers effects.

**Fig. 5:**
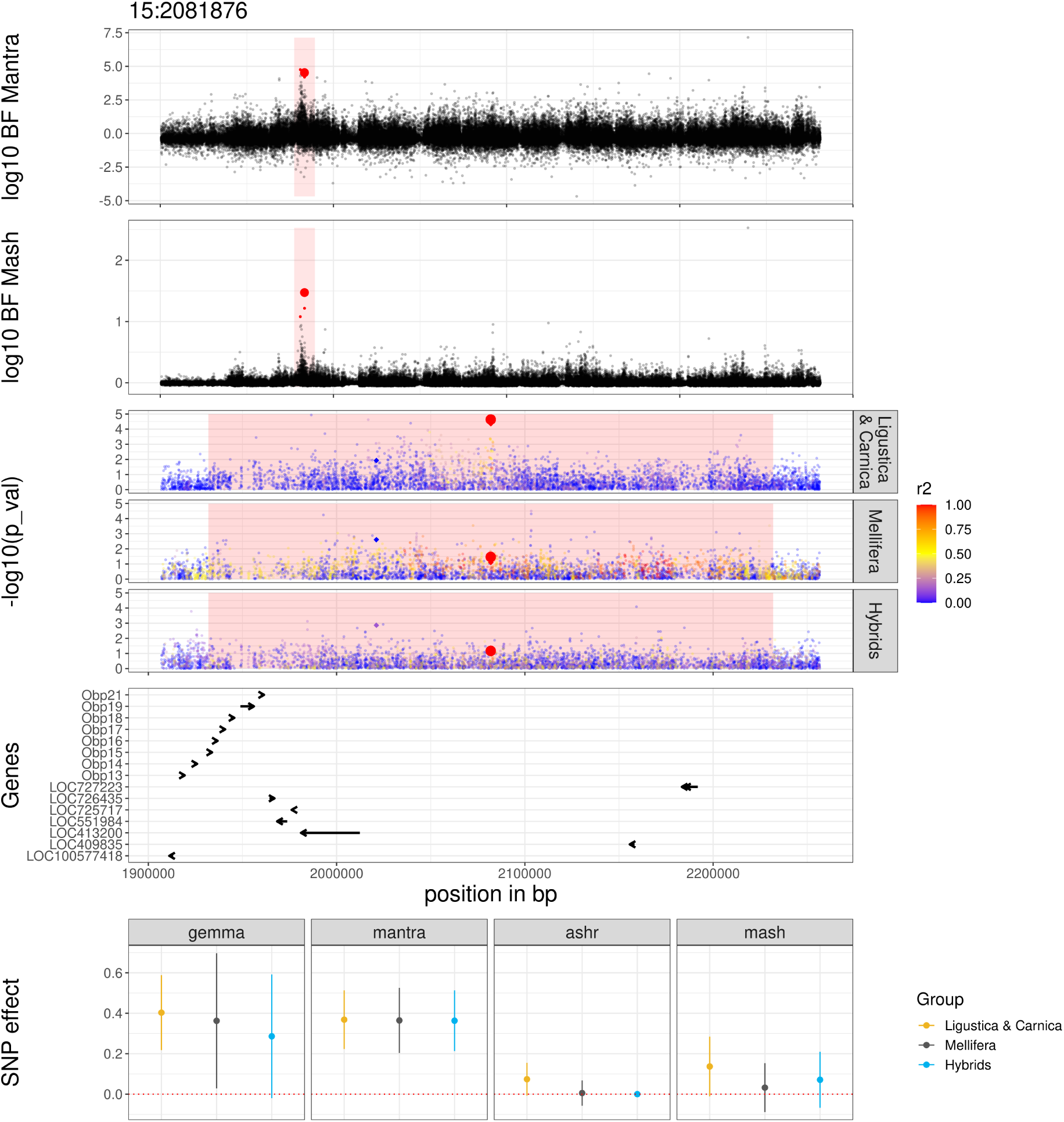
Region on chromosome 15. We plotted manhattan plots for the marker positioned at 2,081,876 bp on chromosome 15 and the surrounding 0.3Mbp region. The marker 2,081,876 is represented by a large red dot when the close by markers identified as near significant thresholds are represented as smaller. From top to bottom we plotted log10(bayes factor) for MANTRA, then for mash, then −log10(p-values), estimated using GEMMA, for the groups *A. m. ligustica & carnica*, *A. m. mellifera* and hybrids. Each marker is coloured based on its linkage disequilibrium r^2^ with the marker of interest. The genes falling in the region of interest are plotted and then the effects of the significant marker extracted from genome wide association studies ran with GEMMA, ash, MANTRA and mash.

### Polygenic architecture of varroa resistance

The program GEMMA [43], used to perform individual GWAS, provides estimates for the proportion of phenotypic variance explained by the SNPs (*pve*) and proportion of genetic variance explained by the sparse effects (*pge*) for each trait in each group (Tab. 1 and Fig. 6). PVEs ranged between 0.20 and 0.82 (se=[0.09; 0.19]). PVEs, estimated using the Bayesian Sparse linear mixed model (BSLMM-GWA), were close to the lmm estimates (LMM-GWA), and ranged between 0.14 and 0.82 (se=[0.07; 0.15]). The 95% confidence and credible intervals for PVEs from both LMM-GWA and BSLMM-GWA appeared exclusively positive, except for *MNR* in the hybrids group. However, PGEs were much lower, ranging between 0.06 to 0.33 (se=[0.08; 0.25]). Their 95% confidence/credible intervals often included zero. The only traits and groups having PGEs different from zero were *MNR* and *recap* for the group of colonies of *A. m. ligustica* type. The estimates for the GWAS on hybrids and *A. m. mellifera* always showed larger standard error, as expected due to the smaller sample sizes of these groups. Interestingly it appears that PVE estimate is slightly higher for *A. m. mellifera* for the *MNR* phenotype compared to the two other groups, whereas they seem similar between the three groups for the two other phenotypes, *varroa_inf* and *recap* (a complete summary can be found in supplementary table SR8).

**Fig. 6:**
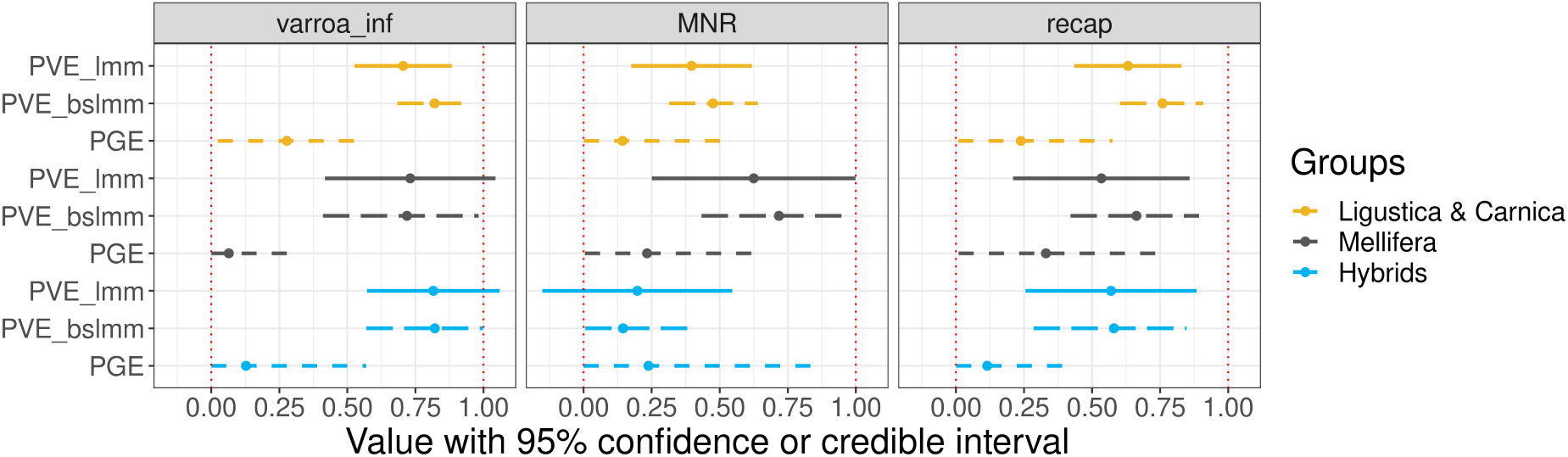
Genome wide association estimates. Confidence and credible intervals for percentage of variance explained by linear mixed model (full line), Bayesian sparse linear mixed model (long dotted line) and percentage of genetic variance explained by Bayesian sparse linear mixed model (short dotted line) for the three group analysed, in yellow *Apis mellifera ligustica & carnica*, in dark grey for *Apis mellifera mellifera* and in blue for hybrid.

**Tab. 1:**
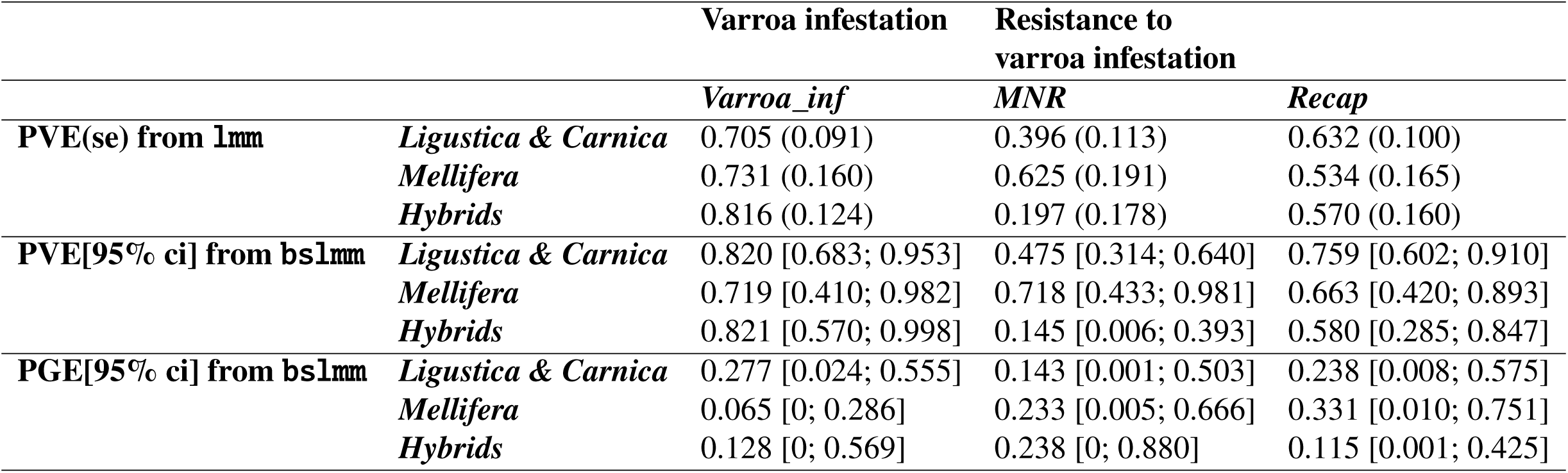
Summary of PVE (Phenotypic Variance Explained) and PGE (Proportion of Genetic variance Explained).

Correlation between SNP effects, across the whole genome, estimated using GEMMA and ash, for individual GWAS analysis, on the one hand and MANTRA and mash, for meta-GWAS, on the other hand were positive. However, as expected, between the different groups there was no correlation when estimated from individual GWAS whereas there was some positive correlation when estimated from meta-GWAS. Across the different phenotypes, *recap* and *MNR* appear to be slightly positively genetically correlated, as one could expect knowing that recapping behaviour, performed by adult bees, is potentially a component of the *MNR* phenotype. The phenotypes *varroa_inf* and *MNR* were not genetically correlated. Finally, *varroa_inf* and *recap* were mostly negatively correlated for *A. m. mellifera* and *A. m. ligustica & carnica* whereas there was no to very little positive genetic correlation between these two phenotypes in the hybrids group (Tab. 2, the detailed correlations can be found in Supplementary table SR5).

**Tab. 2:** Genetic correlations. Range of the estimates from Pearson correlations on the different GWAS analyses methods used in our study.

|  |  | *Varroa_inf* | *MNR* | *Recap* |
| --- | --- | --- | --- | --- |
|  | *Varroa_inf* | 1 |  |  |
| *Ligustica & Carnica* | *MNR* | [0.042; 0.049] | 1 |  |
| *Mellifera* |  | [-0.023; 0.005] |  |  |
| *Hybrids* |  | [0.050; 0.073] |  |  |
| *Ligustica & Carnica* | *Recap* | [-0.304; -0.213] | [0.131; 0.176] | 1 |
| *Mellifera* |  | [-0.132; -0.058] | [0.088; 0.197] |  |
| *Hybrids* |  | [0.021; 0.173] | [0.021; 0.119] |  |

## Discussion

In this study we performed the largest genome wide association study on the resistance of honey bees to their current biggest biotic threat, the parasite *Varroa destructor*. We combined an extensive genotyping and phenotyping effort with meta-analyses methods to identify genetic markers and associated genes harbouring a significant effect on varroa resistance. This leveraged both multiple traits associated to varroa resistance and the complex genetic structure found in honey bee colonies. Our results show that we were able to pinpoint significant effects of some regions of the honey bee genome, within specific genetic types and across the whole meta-population, offering insights into the biological mechanisms involved in varroa resistance. These genomic regions only explain a small portion of the genetic variation which remains mostly polygenic. However we found the contribution of genetics to varroa resistance to be substantial, offering positive perspectives to a possible adaptation and selection of honey bee populations to this relatively recent threat.

Based on 1,500 sampled honey bee colonies, sequenced in pool and genotyped for around 3 million SNPs, this study benefits from the largest sample size (phenotyped and sequenced) known thus far to perform genome wide association study in honey bees: it is the most global association study for honey bee varroa resistance traits to date. The set of colonies studied is highly representative of many honey bee populations worldwide, measured in terms of the genetic backgrounds described in the diversity panel of Wragg et al. (2022) [40]. This study stands out from the previous honey bee quantitative genetic studies that mostly focused on specific genetic backgrounds, often in small experiments, not representative of the field situation and for really specific phenotypes ([38] for a review). In addition to the experimental effort to gather the raw data, this analysis benefited from dedicated statistical methods for reconstructing the honey bee queen genotype from a pool sequencing experiment [31] and meta-analyses approaches to increase statistical power to detect significant associations [41, 42].

### Phenotyping varroa resistance

Resistance to varroa is a complex trait, involving many different aspects of the biology of honey bee colonies. A recent extensive review of varroa resistance traits [26] reveals that there is no clear evidence for significant correlations between the standard traits measured as proxy for varroa resistance, such as varroa sensitive hygiene (VSH), grooming, mite non reproduction (MNR), hygienic behaviour or uncapping-recapping as cleaning behaviour. Depending on the population on which the trait has been measured, in particular their evolutionary history (*e.g.* natural or artifical selection) the correlations between these traits range from ’apparent link’ to ’no link’.

Here, we observe a slightly positive correlations between traits linked to varroa resistance, *MNR* and *recap* overall or at the population level, and a small negative correlation between these two traits and *varroa_inf* (Fig. 2 and in supplementary figure SM1). This finding fits with the hypothesis that there is a panel of mechanisms allowing honey bee to resist varroa and that these mechanisms do not seem to be completely shared across populations or genetic ancestries.

An important experimental aspect that limits varroa resistance studies, including this one, is that most currently applied measures of varroa resistance are difficult to scale to large samples. For example, they can imply tedious and potentially subjective scoring, induced or artificial varroa infestation, multiple measures in time, estimation of ratios and applying heuristic thresholds for minimum detection. However, direct estimates of varroa infestation are the most simple traits to measure to summarise varroa resistance. Indeed, a low varroa infestation can be explained by multiple phenomena either linked to the environment, beekeeping practices, varroa biology or an action from the honey bee colony itself. In this study, in addition to the classical measures of varroa infestation, we proposed to measure varroa infestation indirectly, using the ratio of reads mapped to the varroa mitochondrial DNA over those mapped to the honey bee genome (*varroa_mito*). We believe this new measure offers specific advantages for the study of varroa resistance: (i) there is a high correlation between this estimate and the phoretic varroa infestation (supplementary methods 1), a trait that is more complex to measure, (ii) both colony sequence and varroa infestation information come from a unique biological sample, (iii) there is no potential bias due to the collector, (iv) no specific technical skills are needed on the field and (v) it is comparable across studies. Using it as a phenotype for varroa infestation would facilitate the establishment of a large collection of standardised phenotypic records, necessary to be able to build up information through time, follow phenotypic progress in a surveyed populations, perform genetic meta-analyses and potentially breed honey bee populations for varroa resistance.

#### Insights into biological mechanisms underlying varroa resistance

Varroa resistance mechanisms can be partitioned into two types of traits: first, traits related to hygiene (including VSH, recapping and MNR, but also more broadly grooming behaviour) that involve the accurate detection by workers of varroa infested cells and second, their subsequent inspection/destruction. It has been shown that VSH bees target more specifically cells with highly compromised brood, which is related to the level of infestation in the cells [47, 14]. As a result, cells with fewer mites or mites that are not effectively reproducing are more likely to stay intact, thus increasing the level of mite non reproduction in the colony (MNR). The second type of trait is a trait expressed by either the workers or the brood, that would disrupt mite reproduction within capped cells (and thus increase MNR). Both trait types can reduce mite infestation in the colony, thus increasing varroa resistance of honey bee colonies. Interestingly, in this study we found markers associated with genes that relate to these two categories.

##### Impairment of mite reproduction

We took advantage of the recent review by Mondet et al. (2020) [16] that summarises the results from previous association studies on varroa resistance. We performed a liftover to obtain markers and regions coordinates over the latest genome assembly (HAv3.1, [45]) (supplementary table SM3). Two of our significant SNPs, for *recap* fell into a region around 7Mb on chromosome 1 described as a potential quantitative trait locus (QTL) for VSH, in an association study by Tsuruda et al. (2012) [48]. Two other significant markers 1:2891204:G>A and 1:21374478:G>A fell inside genes LOC410758 and LOC413968 respectively. These genes have been identified in a study by Saelao et al. (2020) [49] looking for selection signal in hygienic honey bee populations of the US. In addition, the marker 11:14369154:G>C, significant for *recap*, is located close to the TpnCi gene, coding for troponin C type. This gene has been shown to be over-expressed in non-hygienic Africanized bee lines, when compared with hygienic lines, by Teixeira et al. (2021) [50]. Finally markers 4:10789077:T>C and 8:1551638:C>T fell into genes found by Ament et al. (2011) [51] as linked to protein abundance in fat body and haemolymph of the adult honey bee. We know that varroa, while infesting the colony, survives by feeding on these bee biological fluids, the fat bodies when on adult bees and the haemolymph when on pupae [52]. These markers might signal either an impact of varroa infestation, or the resilience to such infestation, on biological pathways linked to composition of the honey bee haemolymph and fat body.

In the region on chromosome 8, we identified multiple significant markers close to the gene Ecr, ecdysone receptor. This gene has been identified as crucial for the reproduction of *Varroa destructor*, within the vitellogenin production pathway, and only produced by honey bees. Hence it stands a key in the interaction between honey bees and varroa [53, 35, 54].

##### Detection of varroa infested cells by honey bees

If we look more into the general biology of the honey bee, we saw that two markers, significant for *MNR*, on chromosome 10, fell into the 5-HT2beta gene. This gene is known to be a serotonin receptor, thus involved in olfactory pathways in a large number of insects [55, 56]. The biological hypothesis for its role on the resistance to varroa infestation by honey bee would bee through cues for the adult bees to perform behaviours leading to better colony resistance to the mite. Olfactory biological pathways thus appear crucial to such response to the infestation. We also noticed that the significant markers 3:12973246:A>G and 12973248:A>G fell into LOC413503 (alias GB41230) also found by Mondet et al. (2015) [47], as being differentially expressed in antennae of honey bees expressing VSH behaviour.

In the region on chromosome 15, we identified multiple significant markers located within less than 1 Mb downstream from a group of genes coding for odorant binding proteins. These genes are found in two major clusters on the honey bee genome, seven on chromosome 9, nine on chromosome 15 (monophyletic group called C-minus subfamily); and two on chromosome 10 and one respectively on chromosome 2 and 12, summing to 21 genes [57, 58]. The cluster located on chromosome 15 contains OBPs 13 to 21, 6 of which have already been mentioned in genomic studies looking at varroa resistance related traits (obp 14, 15, 16, 17, 18 and 21; [16, 59]). In particular, Obp14 has been identified as up-regulated in two studies looking at gene and protein expression in VSH bees [47, 60, 61]. Obp18 has been identified by two proteomic studies looking at the VSH and hygienic behaviour traits [62, 61, 60, 63]. Even though these genes were not declared significant in the dedicated analysis we can still hypothesise that they might be involved in some resistance mechanism targeting varroa infestation as they play a major role in sensory pathways. These genes might be relevant for marker assisted selection, as suggested by Guarna et al. (2017) [64] in the case of a tool dedicated to Canadian honey bees selection for hygienic behaviour.

### Genetic architecture of varroa resistance

The review by Guichard et al. (2020) [26] reported heritabilities ranging from close to 0 to up to 0.85, with large standard errors, for traits associated with resistance to varroa. More recently Gabel et al. (2023) [65] estimated the heritability of *MNR* to be close to 0.4, which is in the same range as our estimates. Most 95% confidence/credible intervals for heritability estimates found in literature included zero, while those estimates that did not were mostly modest (< 0.2). In addition, some repeatability estimates of these traits are low (e.g. [25, 21]). More importantly, estimates based on different populations, e.g. *A. m. mellifera* or *A. m. ligustica & carnica* show discrepancy. This could be explained by different genetic architectures involved in these traits.

The heritability estimates from this study, for honey bee resistance traits, seem high compared to standard traits measured on livestock species, which could be due to not being able to fully disentangle genetic from environmental stratification. When intending to estimate heritabilities in honey bee one faces the challenge to integrate the potentially large impact of environmental variation. Part of the population structure is possibly associated with such variation, in addition to genetic variation. In this study, we aim to correct for such structure in our sample by thoroughly accounting for covariates, principal components of the genomic relationship matrix. We computed correlations (using Mantel test) between genetic, geographic and environmental distances between colonies and found that environmental variables are slightly correlated with population structure, whereas their correlation with our phenotypes of interest is not significantly different from zero (supplementary table SR9). Environmental variables seemed to have some impacts that we do not take into account in our analysis (supplementary table SR10), and that might have affected slightly our heritability estimates (upward bias). We can measure the extent of this confounding by comparing the GWAS results obtained by performing single locus GWAS (LMM-GWA) or multi-loci GWAS (BSLMM-GWA), because the former corrects for confounding using principal components of the GRM while the latter does not. Consistent with a small effect of phenotype / genotype confounding, we found the PVE estimates with BSLMM-GWA to be usually larger than those obtained with LMM-GWA (Fig. 6, Tab. 1) but the difference was always very small. Overall, we cannot rule out some inflation of PVE estimates due to remaining confounding but it is not likely to affect our general.

The proportion of genetic variance explained by large effects (PGE), estimated with the BSLMM-GWA [43] was generally low and included 0, except for *varroa_inf* in *A. m. ligustica & carnica*. These estimates support our hypothesis that varroa resistance traits are highly polygenic and not simply driven by a few markers with large effects. The traits linked to varroa infestation and resistance seem to have a small yet significant part of genetic heritability, and thus can be passed on from one generation to next through selection. This is consistent with the few examples of the efficiency of artificial selection for honey bee resistance [14, 24]. Our results obtained in more diverse populations imply that genetic selection, natural or artificial, has the potential to drive increased resistance in other contexts, a positive perspective for honey bee populations worldwide. However, and even though we identified genetic markers with significant effects, it is unlikely that large causal mutations, explaining a big part of the phenotypic variance can contribute significantly to this adaptive response.

### The future of genome-wide association studies in honey bees

The honey bee genome harbours some specificities. First, it is known to experience a large number of recombination events, with an average recombination rate of 37cM/Mbp [45]. Secondly, the effective population size (i.e. the number of actively reproducing individuals) of a local population, or represented in a particular colony, is expected to be rather large, due to the polyandric nature of the natural honey bee mating system, with each queen mating with multiple males from numerous bee colonies within the local mating area, thus avoiding over-representation of a specific animal or genetic line. Such particularities cause low linkage disequilibrium, as seen in the supplementary methods 2, between genomic markers, making it harder to identify candidate loci (QTLs) linked to specific traits and to further select for them.

In addition, the honey bee population exhibits a complex genetic diversity. In this study we provide a better understanding of the genetic background behind varroa infestation and resistance in honey bees in general and the French population in particular. Many honey bee colonies are hybrids of various proportions from the three main *Apis mellifera* subspecies found in Europe, *ligustica & carnica*, *mellifera* and *caucasia*. In this study we took advantage of this admixed population to identify genetic markers linked with our traits of interest within genetic types, in hybrids and across these populations, making it possible to see differences in significance and effect depending on the genetic type. Knowledge of linkage disequilibrium (LD) and associated estimation of local haplotypes for regions of interest combined with knowledge on SNP effects for each sub-species can increase our prediction accuracy for different traits. A better understanding of the local genetic background of the hybrid population could help predict effects of specific SNPs. Studies focusing on hybrid colonies, describing their genetic background throughout the genome and comparing different genetic make up could be highly valuable to identify relevant genetic patterns, especially in the context of genomic selection. As an example, multiple genome regions were flagged with more than one significant marker for the trait *recap* but evidence linking these regions to honey bee biology is lacking. In addition, we identified some markers with opposing effects across the different genetic backgrounds, especially *A. m. ligustica & carnica* and *A. m. mellifera*. It appears relevant to short-list these regions as potential regions of interest for future studies geared towards a better honey bee genome annotation and understanding of underlying biological pathways.

#### Selection on honey bee resistance to varroa

One practical perspective of our work would be to integrate the identified variants into genomic selection program aiming to breed for resistant honey bee colonies. Genomic selection is commonly used in mainstream livestock species. In some countries, such as Germany, beekeepers are grouped in breeding organisations, that make extensive use of artificial insemination to track their mating making the construction of breeding schemes easier [66]. In this context, some studies [67, 68, 69, 70, 71] described the statistical models and sampling strategies that can be successfully applied to implement genomic selection in honey bees. One limitation to the widespread use of these methods is that so far genomic selection has been proven successful when the focus is on a unique honey bee subspecies, e.g *A. m. carnica*, and concentrate their efforts on phenotypes linked to production. In the context of the French honey bee population, as described in this study, the hybrid stock as well as the complex phenotypes of interest, make it less straightforward to directly apply genomic selection. Highly polygenic traits come with challenges when considering selection and we expect that the markers identified in this study will not be sufficient to contribute alone to the establishment of a genomic selection scheme even though the abundant estimated variances for the phenotypes still supports the possibility of selection, as it has been pursued in the US [18]. For future studies, a primary focus should be put on increasing the sample size, in terms of number of phenotyped and genotyped colonies, to boost the precision, detection power and replication capacity of association studies. A way to improve the robustness of our marker contributions to selection decisions would be, for example, to deepen our information on pedigree [71], as well as having access to a large number of individual queen genotypes [72], rather than reconstructed ones. Note however, that accessing this information comes today with a greater experimental burden, potentially limiting study sample sizes. Hence, the right balance to optimize statistical power needs to be evaluated further. In addition, there is a need for standardised biological samples in terms of genotype and phenotype. The genotypes could either come from large SNP panels, covering the whole genome [73, 71] or come from whole genome sequencing experiments characterised using a genetic diversity panel [40]. The phenotypes could be more automated and obtained with reduced sampling variability, as we propose with a novel, sequence-derived infestation measure (*varroa_mito* trait).

Finally, to fully understand the genetic architecture behind varroa resistance one needs to broaden his horizon and look into additional phenomena, not only associated with genetics. For example, it would be necessary to better describe maternal effects, disentangling queen from drones genetic contribution, to our traits.

## Materials and Methods

### Honey bee colonies and sampling strategy

The sampling strategy was established to represent the diversity (both in terms of genetic background and beekeeping practices) of honey bee colonies maintained by French beekeepers. A total of 97 beekeepers, located in France, Switzerland, Luxembourg, the Netherlands, Sweden and New Zealand participated in this study (Fig. 7). Foreign colonies were included as they show similar genetic background to the French honey bee population, mostly because of historical or ongoing trade between beekeepers. A total of 1,513 colonies were sampled to go through sequencing. Out of these 1,513, 1,441 *Apis mellifera* colonies were phenotyped, under the condition that each beekeeper contributed at least 6 colonies (from 6 to 125 with on average 19.25 colonies). These colonies were phenotyped for multiple traits known to be related to varroa resistance (described in details below), once per colony at the end of the beekeeping season (summer and fall in Europe, *i.e.* typically between July and September).

**Fig. 7:**
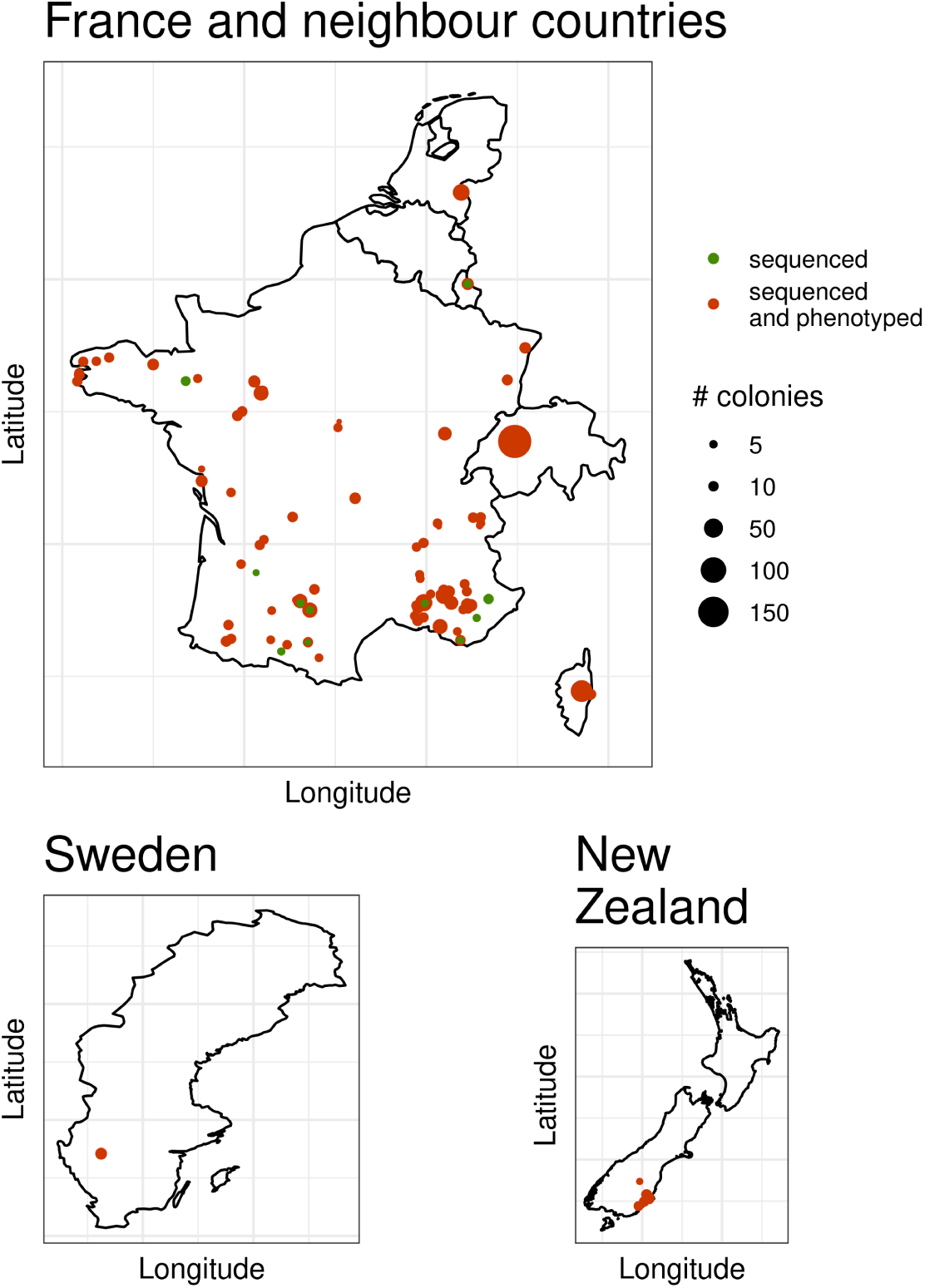
Geographic distribution of the sampled colonies. Geographical locations of colonies that were whole genome sequenced (in green) and both phenotyped and sequenced (in red). The size of the dot represents the number of honey bee colonies per location and category.

### Genetic characterisation of honey bee colonies

All colonies were genotyped from whole genome sequencing of pools of workers using the strategy described in Eynard et al. (2022) [31]. This strategy was decomposed in three steps: first allele counts at Single Nucleotide Polymorphisms (SNPs) were obtained from whole genome sequence, then the genetic background of each colony was estimated from a set of well differentiated markers and finally the genotype of the queen was predicted among colonies of similar genetic background. These different steps are detailed below.

#### Whole Genome Sequencing

For each colony approximately five hundred honey bee workers were ground in 100 mL of TNE buffer. Fifteen mL of ground sample was then collected and centrifuged for 15 min at 3400 rcf (relative centrifugal force). A volume of 200 *μ*L of supernatant was lysed overnight at 56^◦^C, with a solution of proteinase K (Eurobio GEXPRK01-B5) and DTT (1,4 Dithiothreitol). Automated DNA extraction was done with a Qiasymphony^®^-Qiagen. DNA concentrations for each sample were estimated with Infinit200^®^-Tecan. Thereafter, pool sequencing was done on a NovaSeq6000 platform in order to obtain about 30X genome-wide raw sequencing coverage. Sequencing reads were aligned on the honey bee reference genome Amel HAv3.1 (Genbank accession GCA_003254395.2, [45]), using BWA-MEM [74]. In addition, it was expected that the biological sample contained adult varroa, present in phoretic phase on the bees, within the pool of sequenced honey bee workers. The obtained reads were therefore also aligned on the varroa mitochondria (genome Vdes_3.0, Genbank accession GCA_002443255.1, [75]) using the same procedure.

#### SNP Genotyping

Genotypes were estimated at each of the 7,023,976 SNPs identified in Wragg et al. (2022) [40]. Pool sequences were analysed using Samtools mpileup [76] with the recommended parameters: -C minimum mapping quality for reads with excessive mismatches of 50, -q minimum mapping quality for an alignment of 20, -Q minimum base quality of 20. Then Pileup files were interpreted by the PoPoolation2 utility mpileup2sync [77], with a minimum quality of 20 and were finally converted to allele counts and sequencing depth files, filtering out real tri-allelic and potential sequencing error. This procedure led to a set of 6,831,074 SNPs that were used in all downstream analyses. Colonies were sequenced on average with 27.4X coverage, each SNP was on average sequenced with 29.9X coverage, as planned during experimental design (Supplementary table SM1).

#### Population Structure

For each colony, we ran the model presented in Eynard et al. (2022) [31] to estimate the genetic background on 48,589 SNPs selected so that they differentiate the three main genetic background of honey bees in Europe [78, 40, 79]: the **C** lineage comprising the lowly differentiated *Apis mellifera ligustica* and *Apis mellifera carnica*, the **M** lineage of Western Europe *Apis mellifera mellifera* and the **O** lineage of Eastern Europe/South-Western Asia *Apis mellifera caucasia*. Specifically, the SNPs were chosen based on the following criteria: (i) there is a maximum of two polymorphic sites within a 100 base pair window, (ii) only one representative marker per linkage disequilibrium block with *r*^2^ higher than 0.8, (iii) the variance between allele frequencies in the three main European genetic background is higher than zero, to allow for population identification and (iv) so that minor allele frequencies (MAF) within the selected markers follows a uniform distribution. The list of selected markers is provided in supplementary table SM2. For step (ii) above, LD was estimated from the reference diversity panel from Wragg et al. (2022) [40] using the plink software version 1.9 [80, 81] with options –r2 –ld-window 100 –ld-window-kb 10 –ld-window-r2 0.8 and a unique marker was selected manually as the median point for each LD block. The Admixture model from Eynard et al. (2022) [31] was used to estimate the genetic background of each colony (*i.e.* the admixture proportions [82] of each of the three genetic backgrounds).

#### Queen genotype reconstruction

Following the procedure described in Eynard et al. (2022) [31] we grouped the colonies based on their genetic background (*A. m. ligustica & carnica*, *A. m. mellifera* and hybrid) to obtain homogeneous populations. Colonies were if they harboured more than 80% of the same genetic background. Colonies that could not be assigned an homogeneous background colonies were assigned to the ’hybrid’ group. Due to a lack of pure *A. m. caucasia* colonies this group was not considered further in the study. Once homogeneous groups are defined it is possible to perform honey bee queen genotype inference using the homogeneous model described in Eynard et al. (2022) [31]. In short, the method is based on the likelihood of the queen genotype, written as

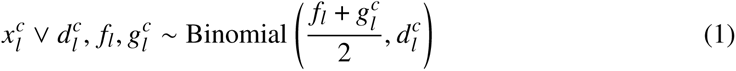

where 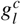 is the (unknown) queen genotype, *f_l_* is the unknown reference allele frequency in the population, 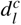 and 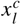 the sequencing depth and allele counts obtained from pool sequencing experiments for locus *l* and colony *c*. By considering all colonies of the same genetic background jointly, *f_l_* can be estimated by maximum likelihood and the posterior probabilities of the three possible genotypes of the queen computed.

### Varroa resistance phenotypes

#### Varroa Infestation

Varroa infestation was quantified with four different measures: phoretic mite infestation (on adult bees using two different methods), brood infestation, and total mite load.

*Phoretic mite infestation* (*v_pho*) was measured using the detergent method [83]. In brief, a sample of approximately 300 adult honey bees was collected in each colony, on a frame containing uncapped brood. After weighing of this sample, the number of mites falling as a consequence of washing with a detergent solution was counted, and the proportion of mites within the sample expressed as the number of varroa per 100 honey bees (assuming the weight of 1 single bee to be 140 mg). An alternative measure of phoretic varroa (*v_mito*) was obtained from the pool sequencing data by calculating the ratio of the number of reads mapping varroa mitochondria on the number of reads mapping the honey bee genome.

*Brood infestation* (*v_brood*) was expressed as the proportion of varroa infesting brood cells in the colony. This proportion was estimated among 300 randomly sampled brood cells on a single frame containing capped brood aged 7 to 11 days post-capping (P5 to P8 stages).

*Total mite infestation* (*v_load*) was estimated by combining the phoretic and brood infestations:

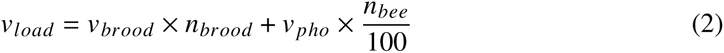

where the total number of brood cells (*n_brood_*) and adult bees *n_bee_* in the colonies were estimated using the ColEval method [84].

#### Recapping of infested cells

The uncapping and further recapping of varroa infested brood cells by adult honey bees is a behavioural trait that has been shown to be associated with varroa resistance [17]. It can be estimated by dissecting brood cells to detect the presence (non-recapped cell) or absence (recapped cell) of the silk cocoon built by the larvae [85]. This was measured on the colony at the same time as measuring mite non reproduction (MNR). This trait is expressed as the proportion of recapped cells within the infested cells.

#### Mite non reproduction

Mite Non Reproduction (MNR), originally known as Suppressed Mite reproduction (SMR), was estimated as detailed in Mondet et al. (2020) [15]. In brief, this estimates infers varroa reproductive status for each brood cell infested by a single varroa foundress and provides a proportion of reproductive mites in the colony. It was estimated on about 300 brood cells (some also used to determine mite brood infestation) with the aim to reach at least 35 single mite infested cells.

All phenotypes were recorded by technicians having followed an extensive training period prior to sampling. Moreover, for statistical analysis they were corrected to fit the assumption of Normality underlying genome wide association study models. Details on the transformations can be found in Supplementary methods 1.

#### Phenotypic Characterisation of colonies

The correlation between varroa-associated phenotypes within and across groups were estimated using the traditional Pearson’s method. A Principal Component Analysis (PCA) was performed using the R package FactoMineR [86], after imputation of the missing phenotypes using the R package MissMDA [87]. Missing phenotypes appeared due to sampling difficulties occurring when performing the scoring in the field. The PCA was used to extract uncorrelated synthetic phenotypes to test for genetic association.

### Genome wide association studies and meta-analyses

#### Genomic relationship matrix

For each group of genetic background identified earlier, *A. m. ligustica & carnica*, *A. m. mellifera* and the hybrids, only SNPs with a MAF above 0.01 and missing rate below 5% were kept. A genomic relationship matrix (GRM) between colonies of the group was estimated on pool sequencing experiment allele frequencies taking SNP linkage disequilibrium into account through the SNP weights produced by LDAK [88], see supplementary methods 2 for more details. Additionally, in order to describe further genetic structure within the group, a PCA on the GRM was performed, using LDAK [89]. The Horn’s parallel analysis [90] was used to decide on the number of principal components kept to explain the variance. 20, 12 and 16 first components were kept from this PCA for *A. m. ligustica & carnica*, *A. m. mellifera* and the hybrids respectively.

#### Genome wide association

Genome wide association studies (GWAS) were performed for three traits: varroa infestation (thereafter called *varrao_inf*), MNR and recapping of varroa infested cells (*recap*). Each GWAS tested the association between the reconstructed queen genotypes and the phenotype using the univariate linear mixed model (lmm) as proposed in GEMMA [43] at each SNP in turn (LMM-GWA), resulting for each SNP in an estimate of its effect and associated standard error, as well as a p-value. In the GWAS, polygenic effects were accounted for with the GRM described above. In addition, further correction was performed by adding the principal components from the PCA on GRM, selected as explained above, as covariates. This was done to correct for the effects of unmeasured confounders with the genetic structure on the phenotype variation (such as apiaries, beekeeper, year … effects). In supplementary methods 2 we illustrate how the structures of the GRMs correlate somewhat to different environmental structures in the data. Association studies were run for all markers initially available, after filtering for minor allele frequency (MAF) above 0.01 and missing rate below 5%. This led to retain a total of 3,084,335; 2,729,072 and 3,185,994 SNPs for the *A. m. ligustica & carnica*, *A. m. mellifera* and hybrid individuals respectively.

To assess the effectiveness of the correction for population structure, the genomic inflation factor *λ_gc_* was estimated as the median of the chi-squared test statistics divided by the expected median of the chi-squared distribution under the null hypothesis. *λ_gc_* ranged between 1.02 and 1.08 for the GWAS on *A. m. ligustica & carnica*, between 0.98 and 1.03 for the GWAS on *A. m. mellifera* and between 0.99 and 1.04 for the GWAS on the hybrid colonies therefore showing really little inflation or deflation of the p-values associated to the tested SNPs (supplementary figures S9).

To call SNPs significant, a False Discovery Rate procedure was applied, using the adaptive shrinkage method [44] as implemented in the ashr R package. Specifically, SNPs with a local false discovery rate (lfdr) and local false sign rate (lfsr) < 0.1 were deemed significant. The lfdr is the probability, knowing the observed data, that an effect is declared significant erroneously and lfsr is the probability, knowing observed data, that the sign of an effect declared significant is wrong [44]. The proportion of phenotypic variance explained by the SNPs (*pve*), and its standard error, was estimated by the univariate linear model and, using the Bayesian Sparse linear mixed model (bslmm, BSLMM-GWA), with default parameters of 0 t 300 SNPs, 1,000,000 sampling steps and 100,000 burn-in iterations, proposed by GEMMA [91]. This model was fitted with the GRM and associated covariates and we estimated the proportion of genetic variance explained by the sparse effects (*pge*) of the trait as well as 95% credible interval from empirical estimates.

#### Meta-analysis

GWAS results for the same phenotype on the three genetic backgrounds were combined with two meta GWAS methods: (i) MANTRA [41] a meta-analysis method dedicated to combine GWAS results from different genetic ancestries (ii) Mash [42] a general purpose data-driven Bayesian meta-analysis method modelling SNP effects with a mixture of multivariate gaussian distributions with different correlation matrices.

MANTRA and mash were run on all SNPs, using effects (*β*) and associated standard errors estimated with the lmm model of GEMMA. For mash inferences, canonical and data-driven covariance matrices were used. The canonical matrices were estimated automatically by mash. Data-driven matrices were: (i) estimated based on extreme deconvolution from PCA matrices, (ii) based on Fst values between populations, similar to MANTRA and (iii) based on correlation between SNP or gene effects in the different groups. Mash includes an estimation of the residual correlation. In this analysis the simple residual correlation estimation model was preferred, as it outperformed more complex residual correlation estimation models. SNPs with a log10(Bayes Factor) > 5 with MANTRA were called significant, a threshold which was shown to be conservative by Wang et al. (2013) [92]. Mash automatically assigns significance to each marker. In our study the corresponding log10(BF) threshold varied from 1.16 to 1.4 depending on the trait.

Genetic correlations were estimated using Pearson’s correlation coefficients on the allele effects, for each SNP, calculated by our different GWAS methods (individually with GEMMA and ash, and across co-ancestries with MANTRA and mash).

#### Gene prioritisation

The variant effect predictor (VEP) tool from Ensembl [93] was used to identify, based on the honey bee genome annotation, for each of the significant SNPs, its impact on the annotation (stop, gained or lost, missense, frameshift…), its closest genes and their locations (upstream, downstream, intronic region …). Additionally, we identified genes, located elsewhere in the genome in LD regions, containing variants in high LD (*r*^2^ > 0.8) with the significant SNP. Linkage disequilibrium was computed, for each group, using data from the genetic diversity reference panel [40], for each significant SNP with all other variants on the same chromosome using the plink software version 1.9 [80, 81] with options –ld-window-r2 0.8 –r2 inter-chr.

## Acknowledgements

This study was performed thanks to the consent of more than a hundred private beekeepers and beekeeping groups to take part in this project, including Arista Bee Research (NL) and Betta Bees Research (NZ). This study was performed with the support of the LABOGENA DNA team (Yannick Poquet and Christina Sann) and the ITSAP team (Anne-Laure Guirao and Antoine Sudan) for the maintenance of the honey bee colonies from the ITSAP experimental station and Benoît Droz for the maintenance of the Swiss colonies, the multiple technicians employed on the project (Baptiste Biet, Marius Bredon, Samuel Favier, Héléne Gerwis, Coralie Guerry, Albéric Delamotte, Pierre Lamy, Antonin Leclercq, Julie Petit, Diane Roriz, Emilien Rottier, Justine Rougé, Agathe Valette, Chloé Vasseur and Alice Revel) for visiting beekeepers and collecting the data for the project. We thank the sequencing platform GeT-PlaGe Toulouse (France), in particular Tabatha Bulach and Rémy-Félix Serre, a partner of the National Infrastructure France Génomique, thanks to support by the Commissariat aux Grands Invetissements (ANR-10-INBS-0009), for the sequencing. Bioinformatics analyses were performed on the computing facility Genotoul. The authors thank LABOGENA DNA for carrying this project. Thanks to François Gerster for initial discussion and his help to build the consortium and target appropriate funding.

## Funding

MOSAR RT 2015-776 project, FranceAgriMer, the Ministère de l’Agriculture de l’Agroalimentaire et de la Forêt.

BeeStrong PIA P3A, Ministère de l’Agriculture de l’Agroalimentaire et de la Forêt and Investissement d’avenir.

Bundesamt für Landwirtschaft BLW grant no. 627000708, Swiss Federal Office for Agriculture.

## Author contributions

Conceptualization: AV, FM, LG, YLC, BB, BS, FP, AD

Data collection: AV, FM, YLC, BB, BS, AD

Data sampling: FM, BB, BD, MG, the BeeStrong technicians, BL, JM

Laboratory: KT, EL, RM

Sequencing: OB

Data curation: SEE, FM, BB, BD, MG, MN

Methodology: BS, SEE, AV, FG

Investigation: SEE, FM, AV, BS

Visualization: SEE, FM, AV, BS

Results interpretation: SEE, BS, AV, FM, FG, FP, YLC

Supervision: FM, YLC, AD, AV, BS

Project lead: LG

Funding acquisition: AV, FM, LG, YLC, BB, BS, FP, AD

Writing original draft: SEE, FM, AV, BS

Writing review & editing: BB, BL, OB, YLC, BD, AD, LG, MG, JM, FG, MN, FP, KT

## Data and materials availability

Raw sequences are made available under the bioproject accession PRJNA1083455 on NCBI (https://www.ncbi.nlm.nih.gov/bioproject?term=PRJNA1083455&cmd=DetailsSearch). Additional data: count files (from popoolation2), raw phenotypes, accession numbers for the sequences, summary statistics from GWAS analysis and scripts are available in the following repository: https://doi.org/10.57745/HHE4CZ. The scripts to perform the analysis are available on github (https://github.com/seynard/gwas_beestrong)

