## Supplementary methods for "Sequence-based genome-wide association studies reveal the polygenic architecture of *Varroa destructor* resistance in Western honey bees *Apis mellifera*"

Eynard et al.

### 1 Details on phenotypes and statistical transformations applied for association study

#### 1.1 Statistical transformation

In order to fit the assumption of Normality underlying genome wide associations models all our phenotypes of interest were corrected. These phenotypes are of three types: (i) link to varroa infestation, (ii) mite non-reproduction and (iii) recapping behaviour.

##### 1.1.1 Varroa Infestation

Varroa infestation was quantified with four different measures: phoretic infestation (on adult bees, using two different methods), brood infestation, and total mite load.

Phoretic varroa (*v_pho*) was transformed as log(x+1) to allow accounting for 0 values and a transformation using 4th root was applied to varroa mitochondria ratio (*v_mito*). For the brood cell varroa infestation (*v_brood*) we used the posterior mean of a Beta Binomial Model with a uniform distribution as prior, allowing to weigh (shrink) estimations based on the number of infested brood cells which can be quite heterogeneous due to variation in the overall infestation of the colony. A logit transformation was applied thereafter to reduce biased towards low values. Finally, a transformation using 4th root was also applied to varroa load (*v_load*).

##### 1.1.2 Recapping of infested cells

As for the brood cell varroa infestation, and for the same reason of heterogeneity of infestation levels across colonies, the posterior mean of a Beta Binomial Model with a uniform distribution as prior, followed by logit transformation, was used for the recapping estimates (*recap_inf*).

##### 1.1.3 Mite non-reproduction

Finally, for *MNR*, the Empirical Bayes estimate, proposed by Mondet et al. 2020 [1] and Eynard et al. 2020 [2], was used.

##### 1.1.4 Phenotype combination

As the phenotypes linked to varroa infestation were highly correlated they were combined as the first axis of the Principal Component Analysis (PCA) on phenotypes (Figure S1) and thereafter called *varroa_inf*.

Table S1: **Summary of the phenotypes used in this study, the statistical transformations applied and their combination**

| Initial name | Phenotypes description | Transformation | Combination | Final name |
|---|---|---|---|---|
| *v_pho* | Phoretic varroa | log(x+1) | | |
| *v_brood* | Varroa in brood cells | Posterior mean logit | | |
| *v_load* | Colony varroa load | 4th root | PCA | *Varroa_inf* |
| *v_mito* | Depth of sequencing for varroa mitochondrial DNA | 4th root | | |
| *MNR* | Mite non reproduction | Empirical Bayes | none | *MNR* |
| *recap_inf* | Recapping | Posterior mean logit | none | *Recap* |

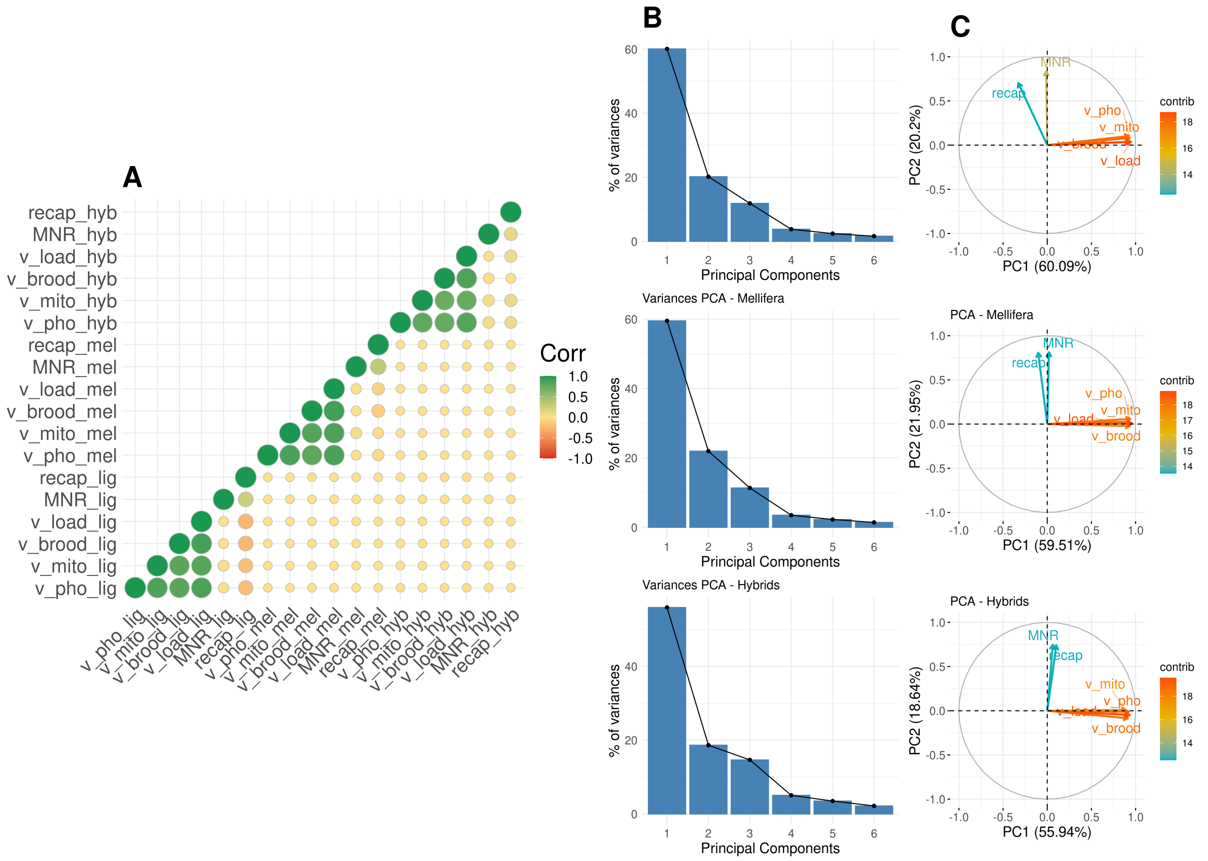


Figure S1: **Correlation and principal component analysis.** Description of the correlation between phenotypes and principal component analysis. (A) gives the correlation between all our phenotypes. (B) summarises the percentage of variance of the principal component analysis explained the axis 1 to 5. (C) representation of our phenotype on principal component analysis for axis 1, 2 and 3, the colour gives the contribution of each variable to the axis, the closer to red, the higher. The correlation and PCA estimates are detailed for each of the three groups defined in our study.

After transformation, the three main phenotypes in each of the groups showed a distribution close to a Normal distribution (Figure S2).

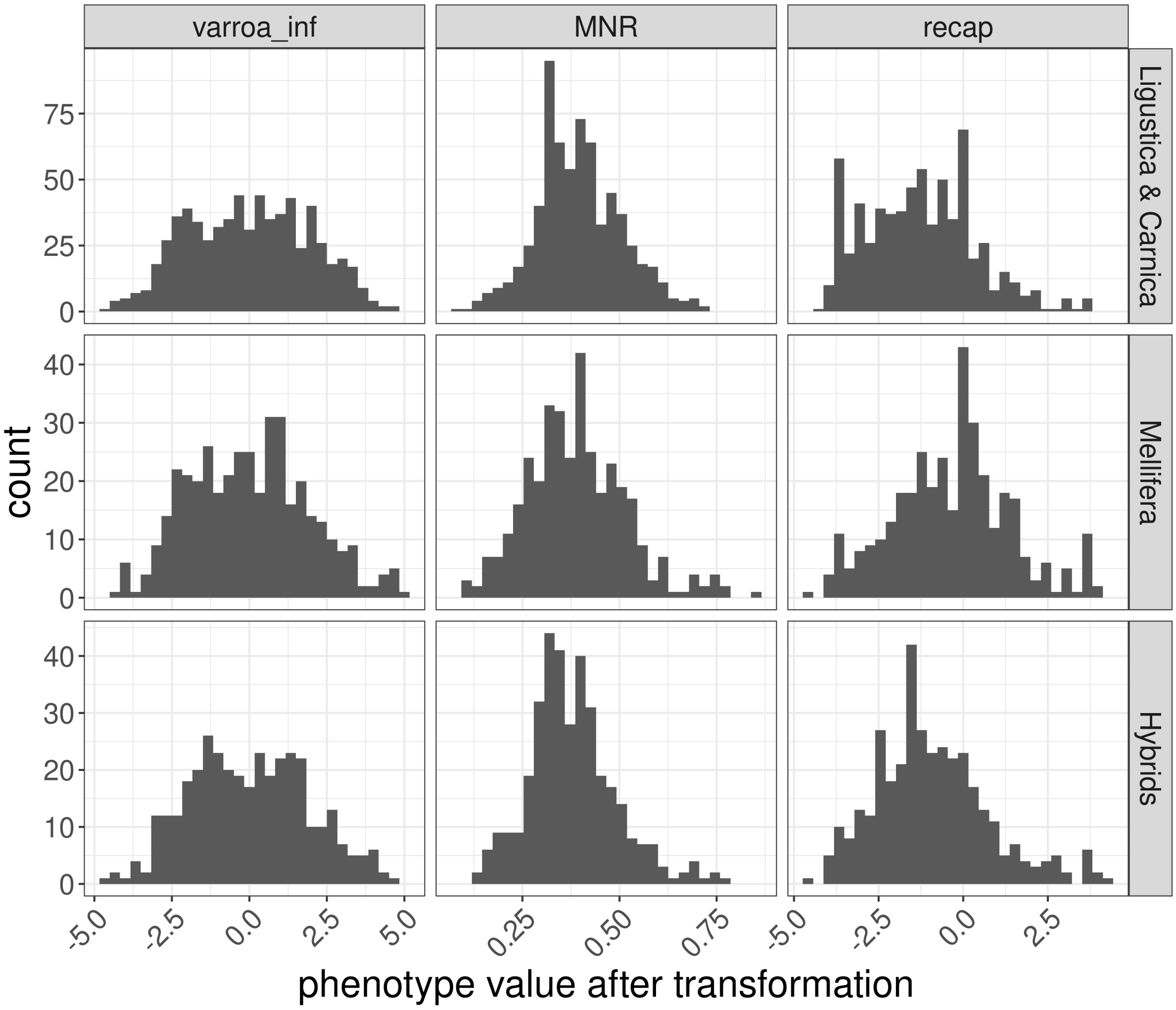


Figure S2: **Distribution of the phenotypes.** Distribution of the three phenotypes used for the Genome wide association study across the three groups.

The number of colonies phenotyped for each of the trait of interest is summarised in Table S2.

Table S2: **Number of samples per group**

| | **Sequenced** | **Varroa infestation** | | | | | **Resistance to varroa** | |
|---|---|---|---|---|---|---|---|---|
| | | ***v_pho*** | ***v_mito*** | ***v_brood*** | ***v_load*** | ***Varroa_inf*** | ***MNR*** | ***Recap*** |
| ***Ligustica & Carnica*** | 703 | 634 | 632 | 667 | 626 | 669 | 667 | 667 |
| ***Mellifera*** | 407 | 397 | 319 | 357 | 351 | 397 | 357 | 357 |
| ***Hybrids*** | 382 | 355 | 332 | 356 | 350 | 357 | 356 | 356 |

#### 1.2 Two methods to measure phoretic infestation

Phoretic infestation was measured in two different ways:

(i) as the count of phoretic varroa for a sample of about 300 adult bees, brought back to a count for 100 bees, following the standard detergent method [3], thereafter called *v_pho*.

(ii) using the depth of varroa mitochondrial DNA sequencing from the pool sequencing experiment, relative to the depth of sequencing of honey bee, as a proxy for the number of varroa found in the adult bee pool sampled, thereafter called *v_mito*.

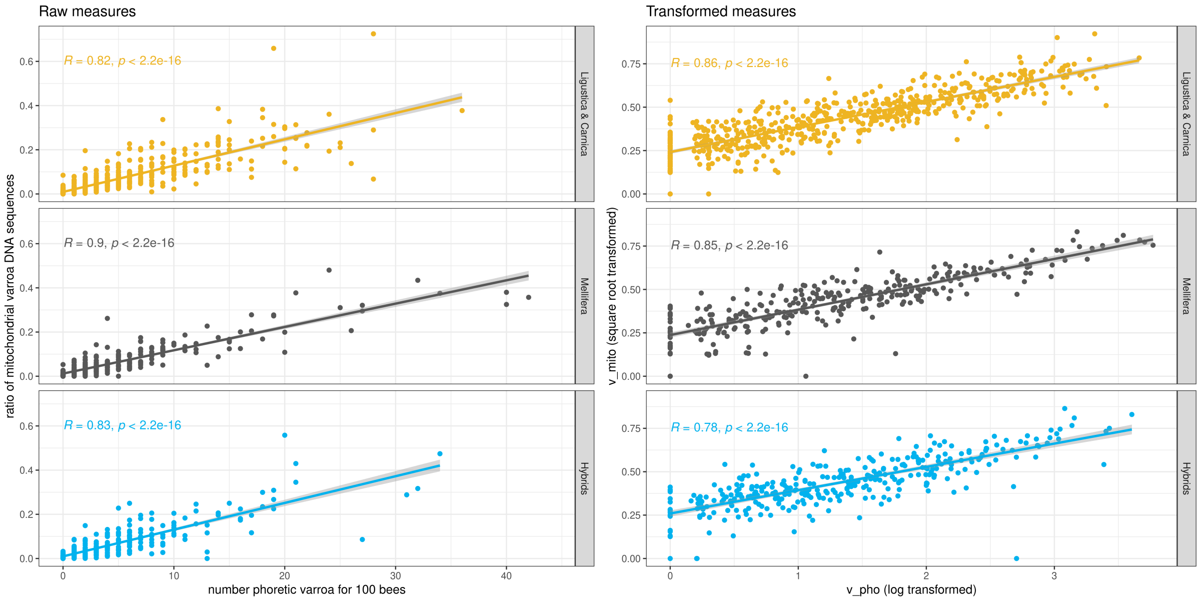


Figure S3: **Correlation between the raw phoretic varroa count and the ratio of varroa mitochondrial DNA and between *v_pho* and *v_mito*, after transformation to follow the normal distribution.** The correlations are estimated for each of the three groups. Correlation coefficient R and its p-value are presented.

We can see that the two estimates for phoretic varroa, raw or transformed, are highly correlated ($R > 0.8$) (Figure S3). It can be noted that for some colonies no phoretic varroa are observed when they are actually measured during the sequencing experiment. Using the sequencing rate of varroa mitochondria seems to be a good alternative to phoretic varroa count. This method would provide with a phenotype in the same experiment as the sequencing, without asking for a dedicated measuring effort from the beekeeper. Unarguably, *v_mito* should be a phenotype of interest for large scale studies focusing on varroa infestation in honey bee colonies.

### 2 Details of the genomic relationship matrix estimations and description of the SNP weights estimated by LDAK

#### 2.1 Genomic relationship matrices

Genomic relationship matrices (GRM) were estimated on data from pool sequencing experiment, thus on allele frequencies. For each of the three groups defined based on genetic background, after filtering for SNPs with minor allele frequency (MAF) above 0.01 and missing rate below 5% we retained 2,832,721; 2,413,514 and 2,912,182 markers for *A. m. ligustica & carnica*, *A. m. mellifera* and the hybrids respectively. GRM estimations, using the program LDAK with weighted SNPs [4], was performed on 292,992; 282,440 and 254,891 SNPs for *A. m. ligustica & carnica*, *A. m. mellifera* and the hybrids respectively, which is about 10% of the set of the total number of SNPs (Table S3).

Table S3: **Number of SNPs used per group after filtering**

| | **Pool seq** | **GRM LDAK** |
|---|---|---|
| ***Ligustica & Carnica*** | 2,832,721 | 292,922 |
| ***Mellifera*** | 2,413,514 | 282,440 |
| ***Hybrids*** | 2,912,182 | 254,891 |

In brief, in LDAK weights are given to SNPs based on linkage disequilibrium S4. Knowing that the honey bee genome is about 250Mbp, that we have about 2.5 million SNPs for this genome and that only 10% of them are kept for GRM estimation after weighing by LDAK we can hypothesise that linkage disequilibrium is lost after 1Kb. Only SNPs having a non-null weight are retained for further GRM estimations. The GRM estimated based on a subset of relevant SNPs seem to better capture the population structure seen in the sample than traditional methods, such as GEMMA [5], when using data like allele frequencies from pool sequencing experiment.

We can observe this pattern when looking at GRM estimated based on allele frequencies or reconstructed queen genotypes for each of our groups or all the colonies together (Figure S5, S6, S7, S8).

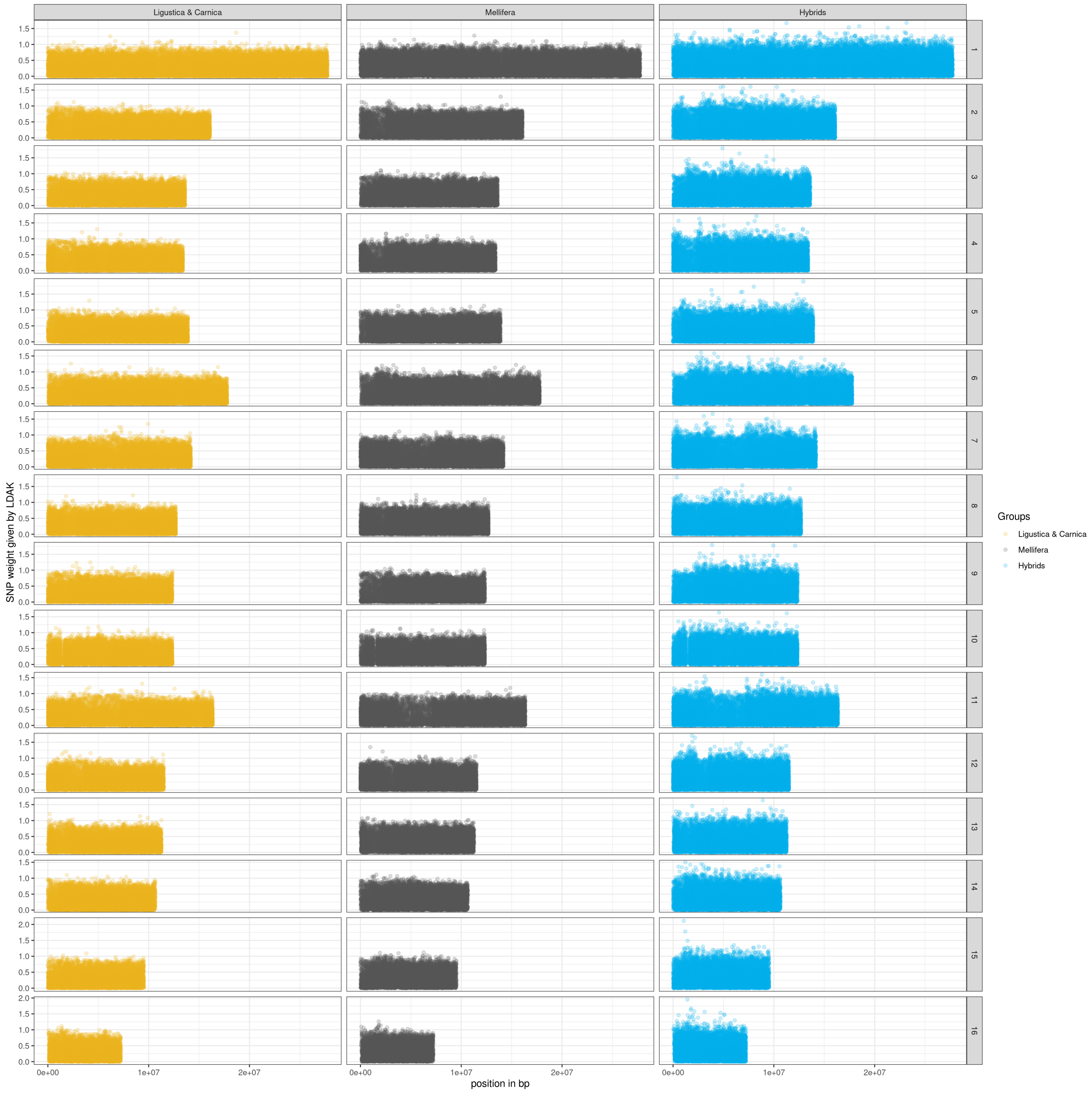


Figure S4: **Values of SNP weights along the genome** Weights given to each SNPs by LDAK, along the genome, for each of the three groups.

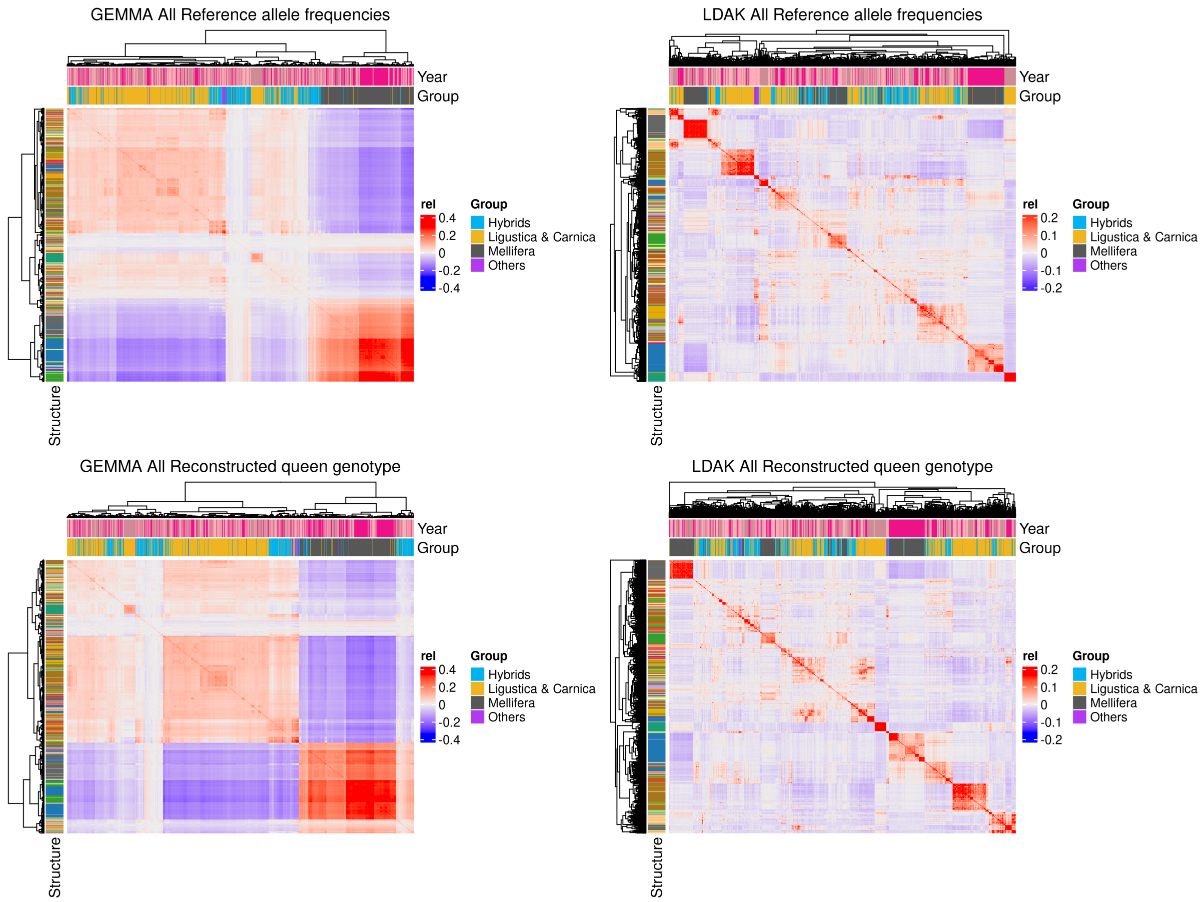


Figure S5: **GRM for all colonies** Genomic relationship matrices estimated from allele frequencies or reconstructed queen genotypes, using LDAK or GEMMA, for all the colonies.

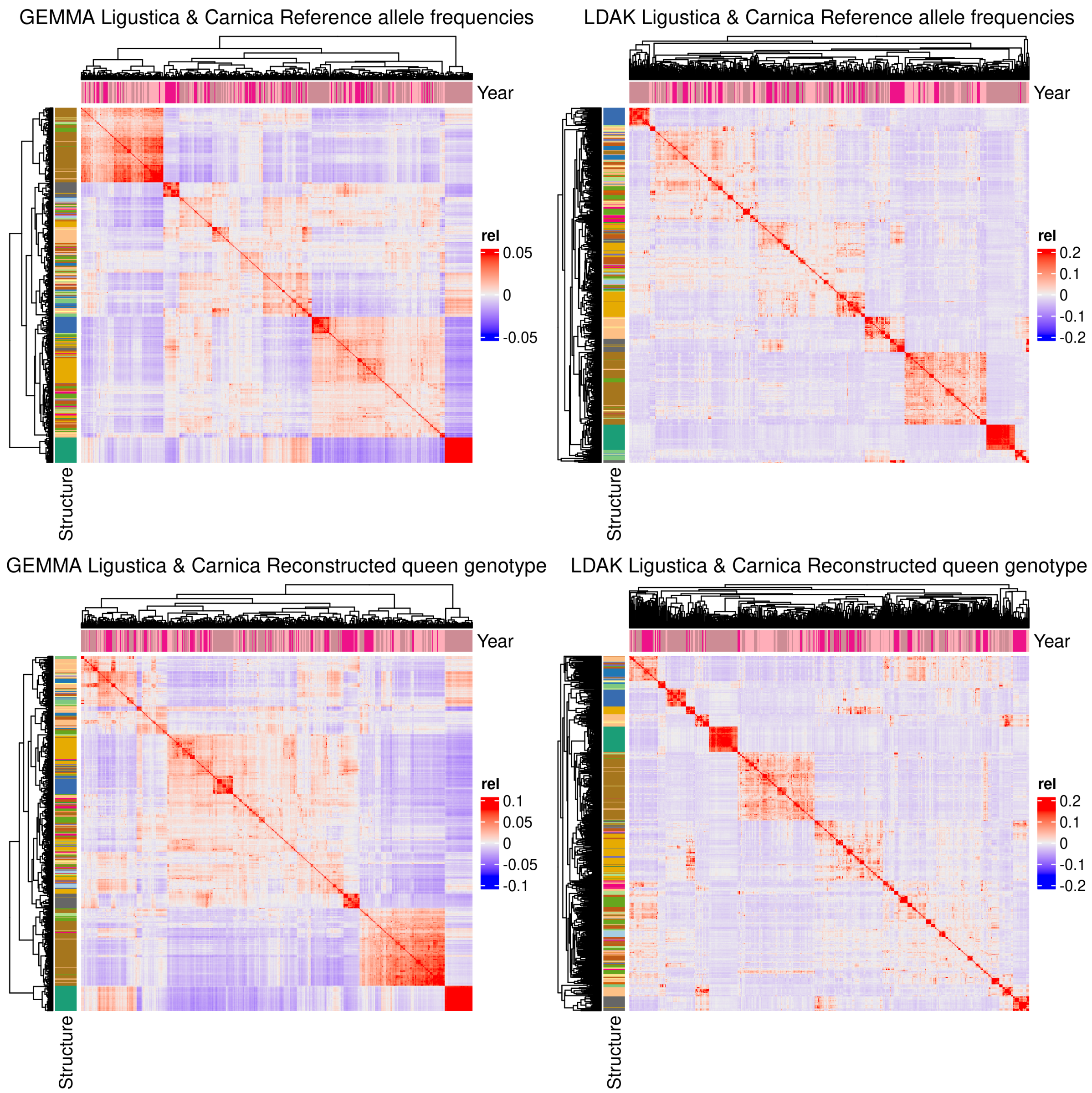


Figure S6: **GRM for Ligustica & Carnica** Genomic relationship matrices estimated from allele frequencies or reconstructed queen genotypes, using LDAK or GEMMA, for the colonies in the ligustica & carnica group.

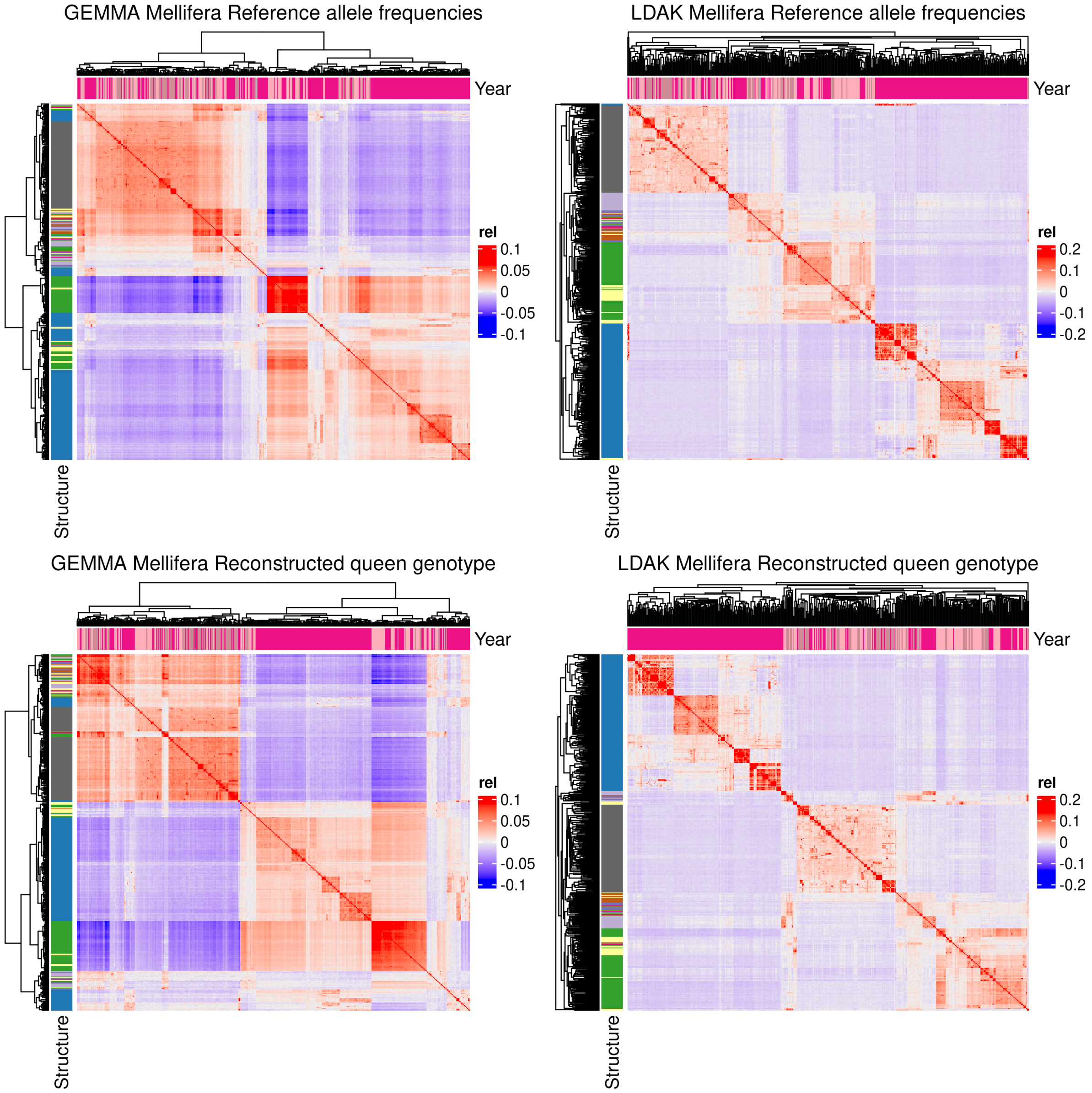


Figure S7: **GRM for Mellifera** Genomic relationship matrices estimated from allele frequencies or reconstructed queen genotypes, using LDAK or GEMMA, for the colonies in the mellifera group.

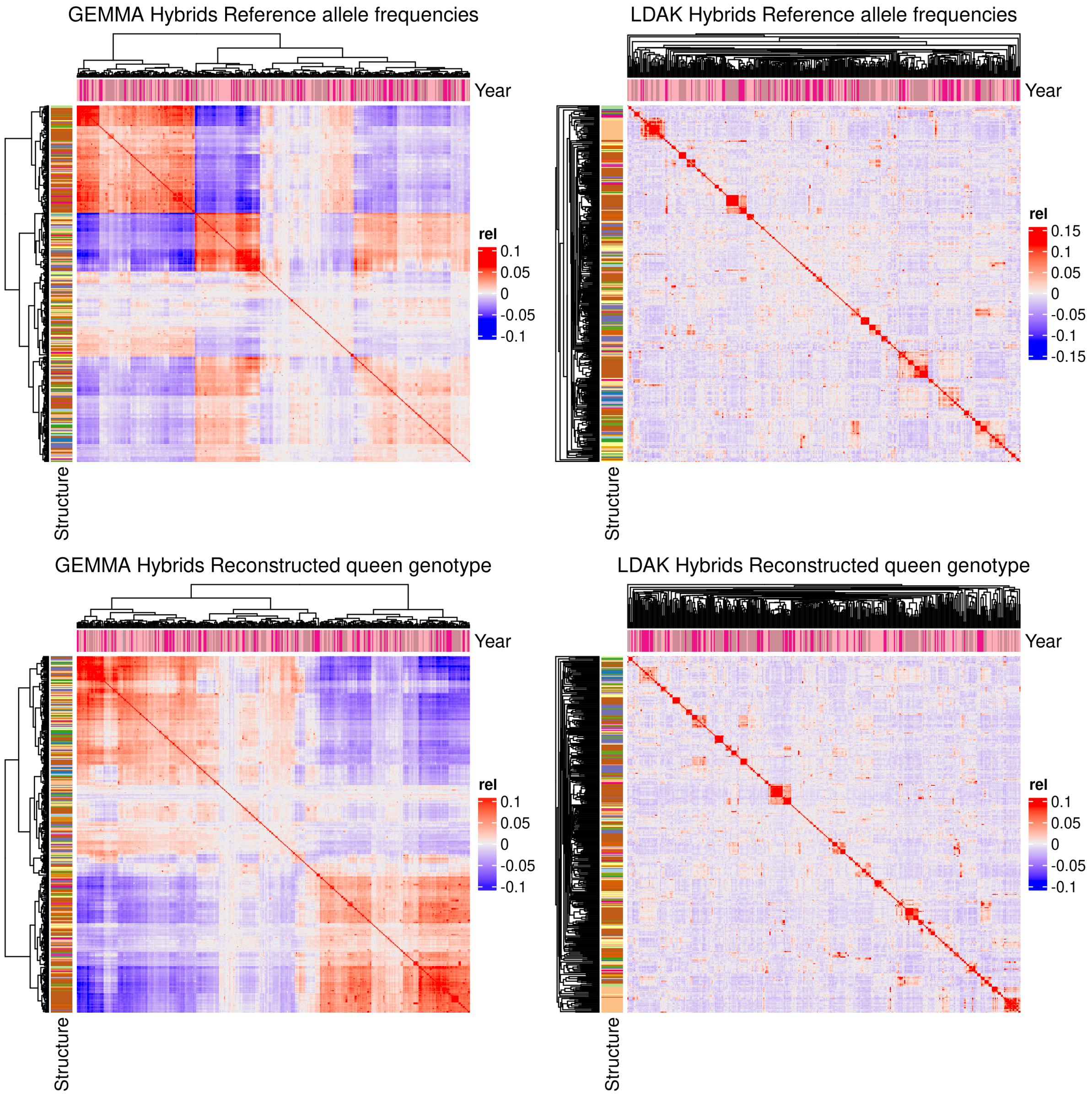


Figure S8: **GRM for hybrids** Genomic relationship matrices estimated from allele frequencies or reconstructed queen genotypes, using LDAK or GEMMA, for the colonies in the hybrid group.

When looking at the heatmaps built from the GRM estimations we can see that in all cases standard estimates, as provided by GEMMA [5], allow to distinguish between genetic background, and large group structures (mostly in correlation with beekeepers association membership), whereas estimations provided by LDAK [4] further discriminate fine structure within each group allowing to better take into account fine within beekeepers association distinction, for example for each individual beekeeper. Using GRM estimated with LDAK allowed to better account for fine population structure in our genome wide association studies (GWAS).

### 3 Supplementary figures

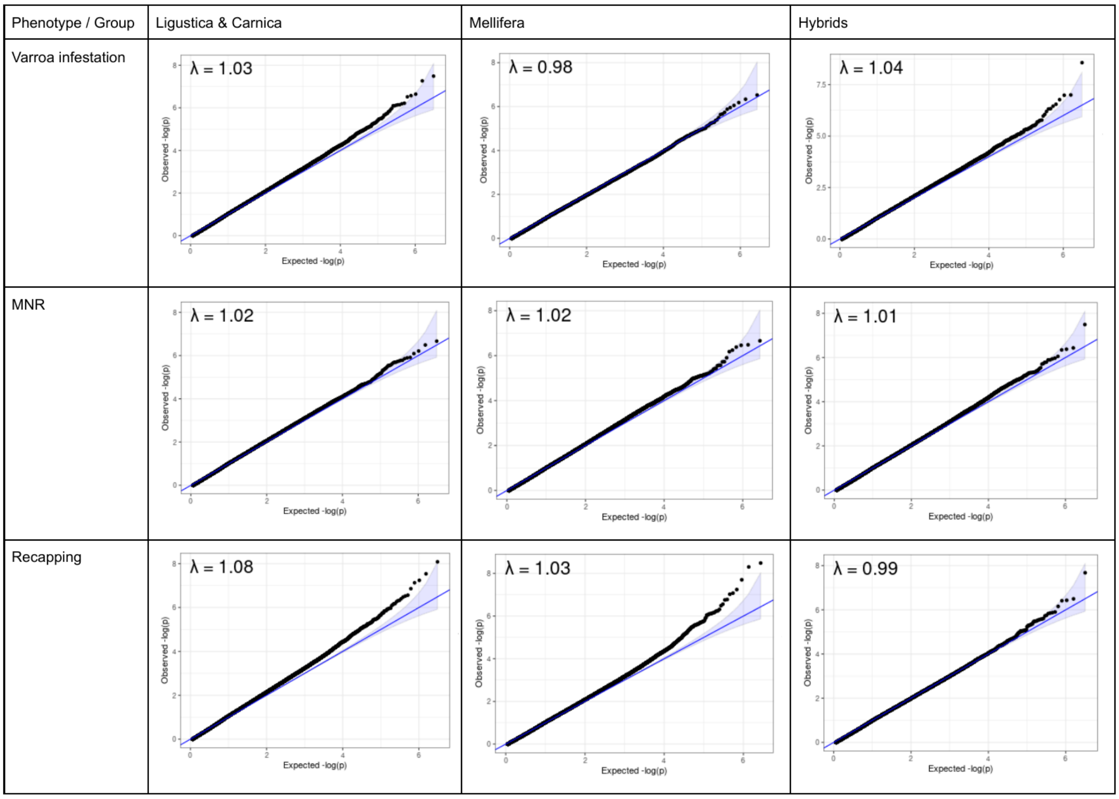


Figure S9: **QQplot for the GWAS on each group and phenotype** QQplot representing the observed -log10(p-values) as function of the expected -log10(p-values) for each of the GWAS performed with GEMMA for the three groups *Apis mellifera ligustica & carnica*, *Apis mellifera mellifera* and hybrids, for the three phenotypes of interest, varroa infestation, MNR and recapping. Lambdas calculated for each of these analysis are reported.

### 4 Supplementary tables (separate file)

**Table SM1**

Detailed sequencing information for each colony (number of SNPs, number of SNPs not sequenced, number of SNPs sequenced, average sequencing depth, minimum sequencing depth, maximum sequencing depth, standard deviation of the sequencing depth).

**Table SM2**

List of the 50k SNPs used for genetic background determination.

**Table SM3**

Liftover of the regions listed in the review by Mondet et al. 2020 [1] from the honeybee reference genome Amel4.5 to the current reference genome AmelHAV3.1. The informations from the review are listed: name, chromosome, start, end of the region strand, size, as well as the information after liftover to AmelHAV3.1: chromosome, start and end of the region, strand, size, coverage, typed, and the character of interest and the associated literature reported in the review.

**Table SR1**

List of the SNPs identified as significant for one of the phenotype of interest in the GWAS per group. Details of the summary statistics for these SNPs are presented: phenotype of interest, group, chromosome, SNP name, position, region (locus), SNP (Reference/Alternative), alternative allele effect, standard error of the effect, p-value, local false sign rate, s-value, local false discovery rate, q-value, posterior mean from ash, posterior standard deviation from ash, closest locus, alias name, variant type from VEP, locus start position, locus stop position, locus strand, SNP position to the closest locus, SNP distance to the closest locus, mention of this SNP/locus in literature, associated phenotype in literature.

**Table SR2**

List of the SNPs identified as significant for one of the phenotype of interest in the meta GWAS per group. Details of the summary statistics for these SNPs are presented: phenotype of interest, chromosome, SNP name, position, region (locus), SNP (Reference/Alternative), number of groups in which the SNP is present, log10 Bayes Factor for MANTRA, posterior probability for MANTRA, number of tested samples, direction of the effect in the different groups, log10 Bayes Factor for mash, number of significant signal in mash, closest locus, alias name, locus start position, locus stop position, locus strand, variant type from VEP, SNP position to the closest locus, SNP distance to the closest locus, mention of this SNP/locus in literature, associated phenotype in literature, comments.

**Table SR3**

Effects of the alternative allele for each of the SNP identified as significant in one or more analysis. In detail: SNP name (chromosome:position), phenotype of interest, study in which it has been found significant, GWAS methods, group, alternative allele effect, standard error of the effect, lower bound of the 95% confidence interval for the allele effect, upper bound of the 95% confidence interval for the allele effect, sign of the allele effect.

**Table SR4**

Region overlap between phenotypes. Are detailed the phenotype combination, the chromosome, the defined region start and stop, the positions of the first, second and if they exist third and fourth SNPs in the region, the loci found in this region, their start, stop position and strand information as well as their biotype.

**Table SR5**

Genetic correlations estimated using a Pearson test across phenotypes within groups and using a Spearman rank test within phenotypes across groups. The details for the correlation estimates, their p-values and the 95% confidence interval for the correlation estimate (in the case of Pearson correlation), the sign of the correlation for each of the GWAS methods, in each group and for each phenotype are presented.

**Table SR6**

List of the SNPs in high LD (r2¿0.8) with SNPs idenfied as significant in at least on of our analysis. In details you can see the chromosome, name of the SNP idenfied as significant, its position, its closest locus, its location relative to its closest locus, the name of SNP in high LD, its position, its closest locus, the r2 value between these 2 SNPs, their distance, the group, the phenotype of interest and the analysis in which the SNP as been found significant.

**Table SR7**

Overlap between haplotype blocks found in Wragg et al. 2022 [6] and SNPs idenfied as significant in our study. We can see the chromosome, the SNP name, its position, the start (BP1) and stop (BP2) position of the haplotype block, the size of the haplotype block, the number of SNPs found in the block by Wragg et al. 2022 [6], in which analysis it has been found significant, the phenotype, the group, the number of genes in the block, their ID.

**Table SR8**

Summary results of the GWAS performed with GEMMA. Are detailed: the sample sizes, the number of PC kept as covariates for GWAS with GEMMA, the lambda values for the QQplot, the PVE and their standard errors, the PGE and their standard errors, the estimate large sens heritability, rho estimates, number of variants with major effects, proportion of variants with non-zero effect (and their 95% confidence intervals).

**Table SR9**

Mantel test performed to compare matrices. Are reported the test statistics, the p-values associated with different tests, and the 95% confidence interval borders as well as the sign of the Mantel correlation.

**Table SR10**

Results for linear regression analysis for each phenotype, each group, to test the effect of environmental co-variates such as the beekeeper, the apiary, the observer ... We report sum of squares, degrees of freedom, F value and p-values.

**Table SR11**

Correlations between allele effects across phenotypes, groups and methods, estimated with Pearson, Spearman rank and linear regression.
